# A matrisome atlas of germ cell development

**DOI:** 10.1101/2023.07.04.547647

**Authors:** Aqilah Amran, Lara Pigatto, Johanna Farley, Rasoul Godini, Roger Pocock, Sandeep Gopal

**Author notes:** Correspondence should be addressed to S.G. and R.P. E mail and. Equally contributed (Aqilah Amran, Lara Pigatto).

## Abstract

The extracellular matrix (matrisome) provides chemical and mechanical cues to control the structure and function of cells and tissues. Yet, comprehensive understanding of how matrisome factors individually and collectively control cell and tissue behavior *in vivo* is lacking. Here, we systematically investigate the function of 443 conserved matrisome-coding genes in controlling germ cell behavior within a complex tissue - the *Caenorhabditis elegans* germline. Using high-content imaging, 3D reconstruction and cell behavior analysis of >3500 germlines and >7 million germ cells, we identify specific matrisome factors that regulate germline structure, protein distribution, germ cell cycle and fate, apoptosis, and oocyte health. These findings reveal matrisome networks acting autonomously and non-autonomously to coordinate germ cell behavior, providing new avenues to study and manipulate cell fates.

## Introduction

The organic extracellular matrix, or matrisome, is a collection of structural proteins, enzymes, growth factors, saccharides, and other small molecules (*1*). The matrisome regulatory output is determined by the independent and collective functions of matrisome components, many of which are post-translationally modified (*2*). Deciphering how the matrisome controls cell fate and behavior requires systematic dissection of matrisome components in a tractable *in vivo* model. Here, we exploit the *Caenorhabditis elegans* germline to investigate how the matrisome controls cell fate and behavior in a complex tissue.

Primordial germ cells are the precursors of mammalian gametes (*3*). The roles of certain matrisome molecules in primordial germ cell development in mammals have previously been described (*4, 5*). Yet, how the entire matrisome collectively controls germ cell fate and behavior *in vivo* is unclear. In *C. elegans*, individual matrisome molecules such as growth factors, structural proteins and proteoglycans can function in germ cell regulation *(6-8)*. However, systematic analysis of matrisome functions in the development of germline stem cells into gametes is lacking. The adult *C. elegans* germline initiates with self-renewing population of mitotic germline stem cells within the distal progenitor zone. As the cells move proximally, they enter meiosis in the transition zone (*9-12*). This is followed by a region of germ cells in meiotic pachytene before they differentiate into sperm and oocytes (*12, 13*). The development of germline stem cells into gametes is controlled by both intrinsic and extracellular signaling (*12, 14-16*). While the role of intrinsic signaling has been extensively studied, the role of extracellular signaling remains unclear (*12*). The ability of adult *C. elegans* hermaphrodites to produce oocytes from undifferentiated germline stem cells provides an avenue to explore matrisome functions at all stages of germ cell development *in vivo* (*12*).

*In silico* characterization of matrisome proteins reveals significant conservation between *C. elegans* and other organisms (*17*). Here, we used RNA-mediated interference (RNAi) to systematically profile the function of 443 conserved matrisome genes in *C. elegans* germline development and germ cell behavior. Our high-throughput analysis of >3500 germlines and >7 million cells surveyed germ cell behavior from immature undifferentiated germline stem cells through to mature gametes. We identified functions for matrisome genes by examining germ cell number, nuclear and plasma membrane morphology, protein distribution, cytoskeletal structure, apoptosis, and oocyte health. By combining experimental and bioinformatic analysis, we identified critical matrisome molecules and their collective, location-specific roles during the formation of gametes from undifferentiated germ cells.

## Results

### Profiling matrisome-associated phenotypes in the *C. elegans* germline

The *C. elegans* matrisome is highly conserved, with 60% of proteins having mammalian orthologs (Fig. S1) (*17*). Gene ontology (GO) analysis shows that *C. elegans* matrisome components have similar biological functions to mammals (Fig. S1). We further categorized the *C. elegans* matrisome into gene families according to established curated databases (Fig. S1). Overrepresentation analysis of comparable mammalian orthologs suggest that *C. elegans* matrisome genes function in 250 signaling pathways, of which 14 pathways have a high confidence level (Fig. S2).

We generated a plasmid library of the 443 matrisome genes conserved in mammals to enable RNAi gene silencing by feeding (Fig. 1A) (*18*). We systematically silenced individual matrisome genes using short-term RNAi for 16 h from the late larval stage 4 (L4) hermaphrodite stage (Fig. 1A). This timing was selected to preclude potential early developmental defects caused by matrisome gene RNAi knockdown and long-term secondary impacts. We used immunofluorescence, 3D reconstruction confocal microscopy, and high-throughput cellular phenotypic analysis to examine germline structure, germ cell number and germ cell cycle, nuclear and protein distribution, plasma membrane structure, apoptosis, and oocyte health in 1- day old animals following RNAi (Fig. 1A). Using this expansive phenotypic profiling approach, we identified 321 matrisome-coding genes that are important for germline development. In our preliminary analysis, we investigated eight phenotypes as numbered - i) progenitor zone cell number, ii) transition zone cell number iii) multinucleated cells in the pachytene region, iv) apoptosis in late pachytene, v) oocyte nuclei morphology, vi) oocyte blebbing, vii) multinucleated oocytes and viii) germ tube morphology (Fig. S3). The number of distinct phenotypes observed per gene knockdown ranged from one to four (Fig. 1B). We observed localized germline defects (212 genes) limited to the distal, pachytene or oocyte regions, and more global defects (109 genes) identified in multiple regions of the germline or in germ tube structure (Figs. 1C, D, S3). A large number matrisome gene knockdowns caused defects in the distal to pachytene germline region, suggesting that these cells are sensitive to matrisome perturbation (Defects i-iv in Fig. 1E). Further, we identified 102 genes important for oocyte development (Defects v-vii in Fig. 1E). A small number of gene knockdowns resulted in defective germline morphology (Defect viii in Fig. 1E). We found that most matrisome gene families have at least one member that controls germ cell development.

**Fig. 1.**
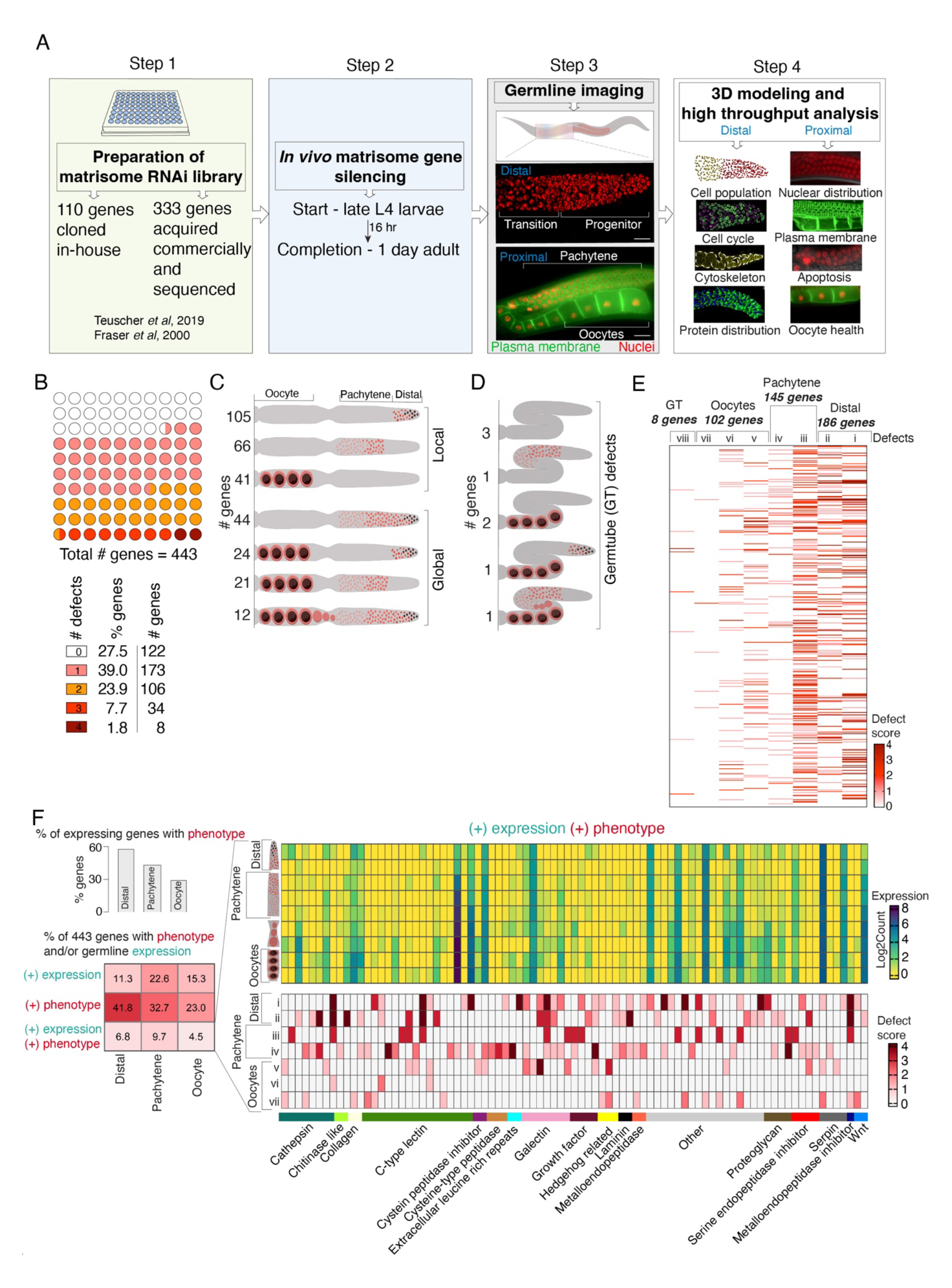
Profiling matrisome functions in the *C. elegans* germline. **(A)** Experimental protocol for short-term RNAi-mediated silencing of matrisome genes and analysis of germline phenotypes. **(B)** Number of defects observed for each matrisome knockdown (0 to 4 phenotypes/per gene). **(C-D)** Location of germline defects following RNAi-mediated gene silencing (distal, pachytene, oocyte regions (C) and/or germ tube (D). Local defects identified at one location, and global defects at multiple germline regions or caused defects in germ tube structure. Number of genes generating the indicated defect after RNAi is shown. **(E)** Heatmap showing the number (bands/lane) and severity (defect score) of phenotypes after matrisome gene silencing. Each band in lanes represents a single gene. Defects are labelled as, i - progenitor zone cell number, ii - transition zone cell number, iii - multinucleated cell at pachytene, iv - apoptosis at late pachytene, v - defective oocyte morphology, vi - oocyte blebbing, vii - multinucleated oocytes, viii - germ tube defects. Defect score shows the severity of the defect. **(F)** Correlation analysis of germline expression pattern and phenotypes of matrisome genes. Bar graph shows percentage of expressed genes at a specific germline region (distal, pachytene or oocyte) that generate RNAi knockdown phenotypes in the same region. Table showing the percentage of total genes with phenotypes and/or expression. Heatmaps show the germline expressed genes that generate at least one phenotype after knockdown in the region of expression. Top heat map – expression level. Bottom heat map – specific defects. Germline schematic shows the location of expression. Defects are numbered as in E.

By examining published expression datasets, we investigated whether matrisome gene expression profiles and germline RNAi phenotypes were correlated (Fig. 1F) (*19*). We quantified the percentage of matrisome genes that are expressed at the same germline location as the phenotype observed following silencing. This revealed 58% (29 out of 50) of genes expressed in the distal germline that when silenced cause distal defects. Similarly, 43% (43 out of 100) of genes expressed in the pachytene region showed pachytene defects, and 29% (20 out of 68) of oocyte expressed genes showed oocyte defects. However, the percentage of genes expressed in the same site as phenotype is relatively low when all 443 conserved matrisome genes are considered (Fig. 1F). We further found that 33 genes that are expressed throughout the germline but phenotypes generated by their silencing were restricted to specific germline regions (Fig. 1F). Finally, we identified 198 matrisome genes that are not germline-expressed yet cause a germline phenotype following knockdown. This suggests non-cell-autonomous germline functions and/or transport of matrisome proteins to the germline from a distal tissue (Fig. 1F). We wondered how many matrisome gene knockdown phenotypes were not previously linked with germ cell/gamete development. We compared our list of matrisome gene hits to genes associated with gene ontology terms ‘germ cell development’ (GO:0007281) and ’gamete generation’ (GO:0007276) in *C. elegans*, *Drosophila,* zebrafish, mice and humans. This identified 15 matrisome genes previously associated with germ cell development and gamete generation, leaving 306 genes not previously associated with these GO terms (Fig. S4).

### Matrisome components control distal germ cell behavior

Germ cell production in *C. elegans* begins at the distal end within the progenitor zone, followed by a closely associated transition zone where germ cells are in early meiosis (*20*). These regions contain defined numbers of germ cells in wild-type animals (*21*). To study the impact of matrisome genes on germ cell behavior, we quantified germ cell number in the progenitor and transition zones following RNAi (Table S1 and S2). We focus here on all matrisome genes except C-type lectins, the largest group of predicted matrisome proteins in *C. elegans*, which will be addressed separately in the manuscript. We found that RNAi silencing of 39 genes affected progenitor zone cell number, 32 genes affected transition zone cell number, and 31 genes affected cell number in both zones (Fig. 2A). Knockdown of all major matrisome gene families, including metalloendopeptidases, cathepsins, serine proteases, and proteoglycans, caused a reduced number of distal germ cells (Fig. S5). The number of cells at the distal germline is maintained by proliferation of germ cells in the progenitor zone. A cohort of germ cells within the progenitor zone act as stem cells that undergo mitotic division to self-renew and generate differentiated gametes (*20*). We performed GO analysis for the mitotic cell cycle (GO:0000278) to identify genes involved in this process in *C. elegans, Drosophila*, zebrafish, mice and humans. Comparison of the genes associated with mitotic cell cycle with our experimentally validated genes revealed that orthologs of *apl-1* (amyloid beta precursor-like protein), *let-756* (fibroblast growth factor), and *ketn-1* (myotilin) are associated with mitotic cell cycle. To investigate any possible indirect links between our experimentally validated genes that regulate progenitor zone number and the mitotic cell cycle, we performed STRING (Search Tool for the Retrieval of Interacting Genes/Proteins) analysis with a combined list of our experimentally validated genes and genes associated with mitotic cell cycle. This analysis provides a probability of experimentally validated genes in regulating the mitotic cell cycle (Fig. 2B). To explore this further, we selected 23 genes that significantly reduced progenitor zone cell number (p <0.0001) to study the germ cell cycle (Fig. S6). Cell proliferation is determined by the time each nucleus spends in S phase (*22*). Therefore, to study the effect of matrisome gene silencing on the cell cycle, we quantified S phase using 5-ethynyl-2′-deoxyuridine (EdU) staining (Fig. 2C) (*23*). First, we established the number of nuclei at each stage of S phase in control germlines (Fig. S5). Using these control values, we calculated the normalized number of nuclei in S phase after each matrisome gene knockdown (Fig. 2D). Silencing *zmp-5* (metalloendopeptidase), *let-756* (fibroblast growth factor), *lec-3* (galectin) and *apl-1* (amyloid ý-precursor like protein) affected the total number of S phase cells (Fig. 2D). Analysis of the early, mid and late S phase revealed additional matrisome genes affecting S phase transitions (Fig. 2E-G). The number of nuclei in all stages of S phase was altered by silencing *apl-1* and *let-756* (Fig. 2D- G), whereas silencing *zmp-3* (metalloendopeptidase) affected only the early and mid-S phase stages (Fig. 2D-G). Finally, silencing *cri-2* (metalloendopeptidase inhibitor)*, W07B8.4* (cathepsin), and *wrt-1* (hedgehog related) altered the early S phase, and silencing *F32H5.1* (cathepsin), *timp-1* (metalloendopeptidase inhibitor), *skpo-2* (peroxidase), and *endu-1* (extracellular nuclease) affected late S phase (Fig. 2D-G). Together, these data confirm two previously identified genes *(apl-1 and let-756)* and reveal 10 matrisome genes that control the mitotic cell cycle (Fig. S5) *(24)*. These results indicate that specific matrisome factors control S phase transitions that likely impact progenitor zone cell number (Fig. 2D-G, S5) (*25*).

**Fig. 2.**
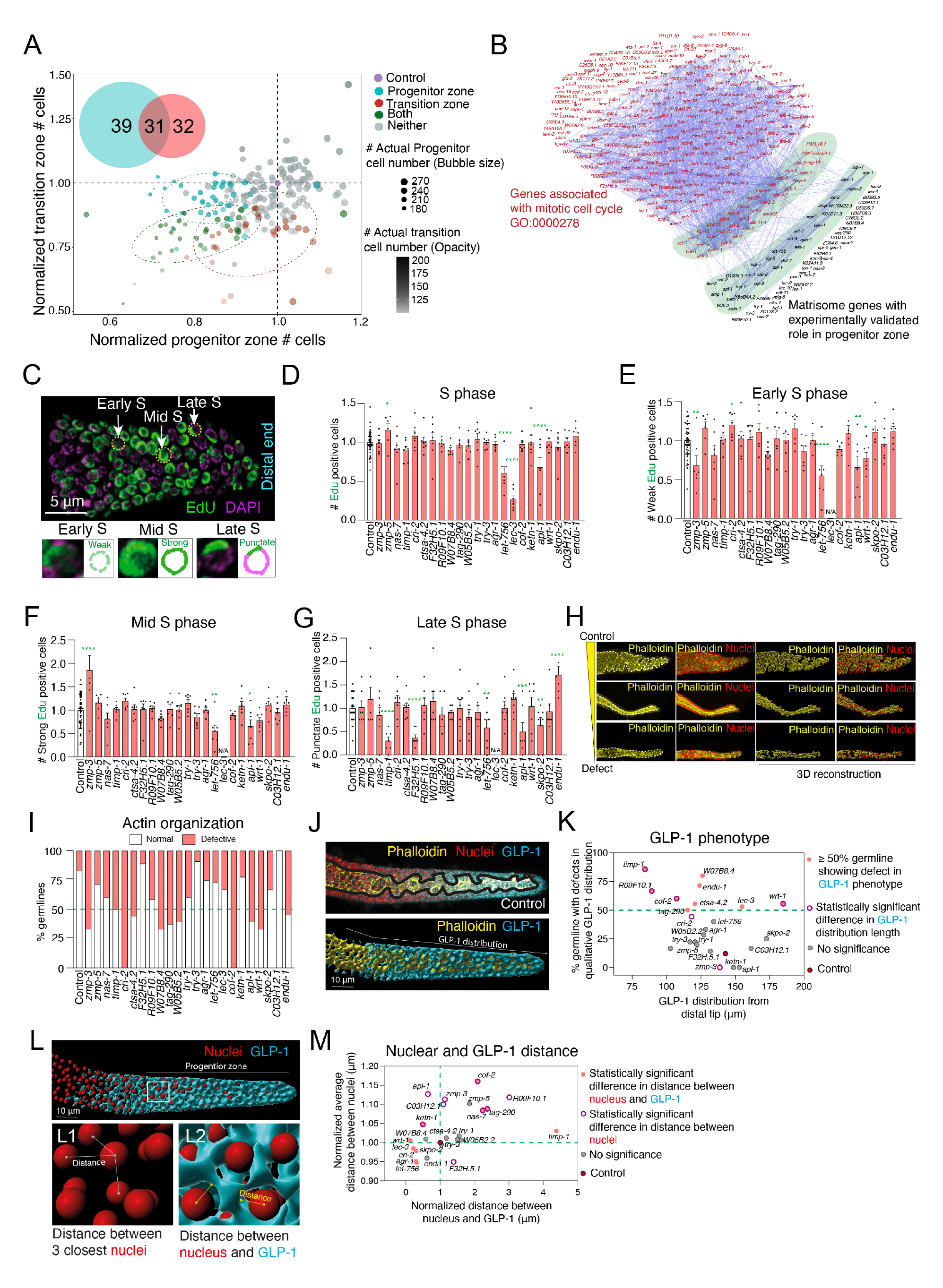
The matrisome controls germ cell number and the distal germline structure. **(A)** Scatter plot showing the number of cells within the progenitor and transition zones after individually silencing all conserved matrisome genes (except C-type lectins). Each circle represents a gene. The position of the circle shows the difference in comparison to control RNAi, which is represented as 1. The size and opacity of each circle is proportional to the number of germ cells at the progenitor and transition zones, respectively. RNAi knockdown of 39 genes affects progenitor zone cells (blue), 31 affects transition zone cells (red), and 30 affects both regions (green). n>7. Blue, red and green = statistically significant. Grey = not significant. **(B)** STRING analysis of association between matrisome genes controlling progenitor zone germ cell number and the mitotic cell cycle. **(C)** Representative images of EdU (green) and DAPI (pink) staining of progenitor zone cells. Intensity and shape of EdU staining indicate S phase stages. **(D-G)** Graphs showing the normalized number of total (D), early (E), mid (F), and late (G) S phase cells following matrisome gene RNAi (red bars) compared to the control (white bar). * p< 0.05, ** p< 0.01, *** p< 0.001, **** p< 0.0001. n>6. **(H)** Micrographs of control (top panel) and defective (middle and bottom panels) germlines showing cytoskeletal structures (phalloidin-yellow) and nuclei (DAPI-red) at the distal end. 3D reconstruction of nuclei positioning on the cytoskeleton. **(I)** Percentage of germlines with normal and defective cytoskeletal structures (corresponding images in Fig. S6). n>7. Gene knockdowns causing ≥50% defects (green line) were identified as significant. **(J)** 3D reconstruction of endogenous GLP-1 (blue) and phalloidin (yellow) in the distal germline. GLP-1 protein distribution and staining phenotype were measured. **(K)** Graph showing the distance of GLP-1 distribution from distal end against the GLP-1 staining phenotype (corresponding images in Fig. S7). Gene knockdowns causing ≥50% defects in GLP-1 phenotype (pink circles above green line) were identified as significant. n = 7. Gene knockdowns causing statistically significant difference in GLP-1 distribution length marked with maroon rings. **(L)** 3D reconstruction of endogenous GLP-1 (blue) and nuclei (red) in the distal germline. Progenitor zone is marked on the panel. L1 and L2 show magnification of marked area. The distances between three neighboring nuclei to each nucleus in progenitor zone shows the nuclear distribution. The shortest distance between GLP-1 and nuclei is quantified for each nucleus. **(M)** Graph showing nuclear distribution (average distance between 3 adjacent nuclei) and GLP-1 proximity (distance between nucleus and nearest GLP-1) in the progenitor zone. n = 7. Statistically significant difference in nuclear distribution and GLP-1 distance marked in maroon rings and pink circles.

An important factor contributing to cell cycle regulation is the access of germ cells to proteins required for cell cycle maintenance (*26*). The germline cytoskeleton maintains a ‘zig-zag’ structure at the distal end that determines germ cell placement and protein distribution, and thus potentially affects their access to proteins necessary for proliferation (Fig. 2H) (*21, 27*). As matrisome molecules play crucial roles in cytoskeletal organization (*28*), we hypothesized that changes in cytoskeletal structure at the distal end of the germline upon matrisome gene knockdown could influence the germ cell cycle. Therefore, we used phalloidin staining to analyze the actin cytoskeletal architecture of the distal germline after silencing the 23 matrisome genes that cause a significant change in progenitor zone cell number (Fig. 2H-I and S7). We detected multiple distinct cytoskeletal changes following matrisome gene knockdown (Fig. 2H-I and S7). Germline defects varied from simple loss of ‘zig-zag’ configuration to total disorganization of the cytoskeleton (Fig. 2H and S7). We found that silencing of 11 matrisome genes (*zmp-3*, *timp-1, cri-2*, *ctsa-4.2, W07B8.4*, *tag-290, W05B2.2, cof-2, apl-1, wrt-1* and *endu-1*) caused defective cytoskeletal structures in at least 50% of germlines analyzed (Fig. 2I, S6). Except for *tag-290*, *W05B2.2,* and *cof-2,* these genes are also required for correct cell cycle progression *(Fig. 2D-G).* To underpin these results, we performed network analysis between the genes with experimentally validated distal germline phenotype and genes associated with the broad gene ontology term ‘cytoskeleton’ (GO:0015629). This identified 42 matrisome genes with experimentally validated defects at the distal end that are associated with the GO:0015629 term (Fig. S7), including five genes (*apl-1, cof-2, cri-2, tag-290*, and *wrt-1*) that have verified cytoskeletal defects (Fig. S7). Together, our results reveal crucial roles for specific matrisome factors in maintaining germline structure that impacts germ cell behavior.

We speculated that the effects of matrisome factors on progenitor zone cell behavior may be caused by changes in distribution of germ cells regulatory proteins due to cytoskeletal defects. To examine this, we studied the GLP-1/Notch Receptor - a critical regulator of germ cell proliferation in the progenitor zone that responds to Notch ligands produced by the distal tip cell *(7)*. GLP-1 protein exhibits a gradient of expression in the distal germline with highest expression at the distal tip (Fig. 2J, S8) (*7*). GLP-1 is also localized around nuclei, forming a boundary (Fig. 2J, S8) (*7, 29*). To study whether changes in germline cytoskeletal structure alters this arrangement, we reconstructed 3D models of distal germlines following DAPI staining and GLP-1 immunofluorescence (Fig. S8). First, we analyzed the distribution of GLP- 1 distance from the distal tip and its placement around germ cell nuclei (Figs. 2J-K, S8). This revealed four genes (*timp-1, R09F10.1, cof-2* and *wrt-1*) that when silenced cause a significant difference in GLP-1 distribution length and ≥50% germlines with defective GLP-1 nuclear placement compared to control. In addition, two genes (*cri-2* and *zmp-3*) showed significant defects in GLP-1 distribution from the distal end and five genes (*tag-290, ctsa-4.2, endu-1, lec-3* and *W07B8.4*) showed altered GLP-1 placement (Figs. 2K, S7). Except for *lec-3* and *R09F10.1*, all these genes also exhibited cytoskeletal defects in ≥50% germlines. Finally, we examined the nuclear distribution and the proximity of nuclei to GLP-1 protein in the progenitor zone (Fig. 2L-M). To study nuclear distribution, we quantified the average distance between each nucleus and its three closest neighbors (Fig. 2L-M). The most significant gene knockdowns (p<0.0001) showed substantial reduction in cell number at progenitor zone. The distance between nuclei would be higher in those knockdowns compared to control if the progenitor zone occupies the same volume. Alternatively, a closer distance suggests densely packed nuclei and a likely shorter progenitor zone. The distribution of nuclei and GLP-1 expression would also determine the proximity of GLP-1 to each nucleus. GLP-1 quantification after RNAi revealed that 8 out 11 genes (*zmp-3, timp-1, cri-2, tag-290, cof-2, apl-1, wrt-1*, and *endu-1*) with a defective germline cytoskeleton had a significant difference in nuclear distribution and/or proximity to GLP-1 protein (Fig. 2M). Together, our analysis of the cell cycle, cytoskeleton, and nuclear/protein distribution suggests that specific matrisome factors control progenitor germ cell behavior through multiple molecular and structural mechanisms (Fig. S8).

### Specific matrisome components control of meiotic cell behavior and oocyte development

As germ cells move distally from the progenitor zone, they enter meiotic pachytene prior to differentiating into gametes. During meiotic pachytene, germ cell nuclei organize within plasma membrane boundaries through cleavage furrow formation and cytokinesis (*21, 30*). Defects in these processes can cause multinucleation of germ cells (MNCs) (Fig. 3A) (*31*). Since matrisome remodeling is thought to play a role in cleavage furrow formation and cytokinesis, we investigated whether matrisome factors control MNC occurrence (*32*). GO annotation using orthologs of *C. elegans* matrisome genes revealed that only annexin and hemicentin were involved in cytokinesis (GO:0000910), cleavage furrow (GO:0032154), cleavage furrow formation (GO:0036089) or cellularization of cleavage furrow (GO:0110070). Using a transgenic strain co-expressing fluorescent markers targeting nuclei (mCherry-histone H2B) and plasma membrane (GFP fusion that binds PI4, 5P_2_), we quantified the number of MNCs per germline (*33*). We found that 11% of control germlines contain MNCs (Fig. S9). We classified a matrisome gene knockdown significant if we observed 55% (five times that of control) or more germlines with MNCs. Using this criterion, we identified 21 matrisome gene knockdowns that increased the penetrance of MNCs (Fig. 3B). This include metalloendopeptidases and their inhibitors (*C48E7.6, acn-1, adt-3, nas-13* and *cri-2*), cathepsins (*asp-4, cpr-2* and *F32H5.1*), growth factors (*tig-2* and *tig-3*), hedgehog ligands (*wrt-9*), galectins (*lec-3, -9* and *-10*), semaphorin (*smp-1*), serine endopeptidase inhibitors (*ced-1* and *ZC84.1*), cysteine rich secretory protein (*scl-19*) and others (*lon-1, mua-3* and *hse-5*) (Fig. 3B). However, short-term RNAi knockdown of annexins (*nex-1, -2, -3 and -4*) and hemicentin (*him-4*) was insufficient to induce MNC, which contrasts a previous report in null mutant animals (*34*). While we identified 21 matrisome genes that prevent MNC, the expressivity of these defects varied (Fig. 3C). For example, silencing *lec-10*, a member of galectin family, resulted in one MNC in every germline. In contrast, *asp-4* RNAi caused up to 12 MNCs in half of the germlines analyzed (Fig. 3C). Taken together, we identified multiple matrisome factors that have previously unknown roles in cytokinesis or cleavage furrow formation.

**Fig. 3.**
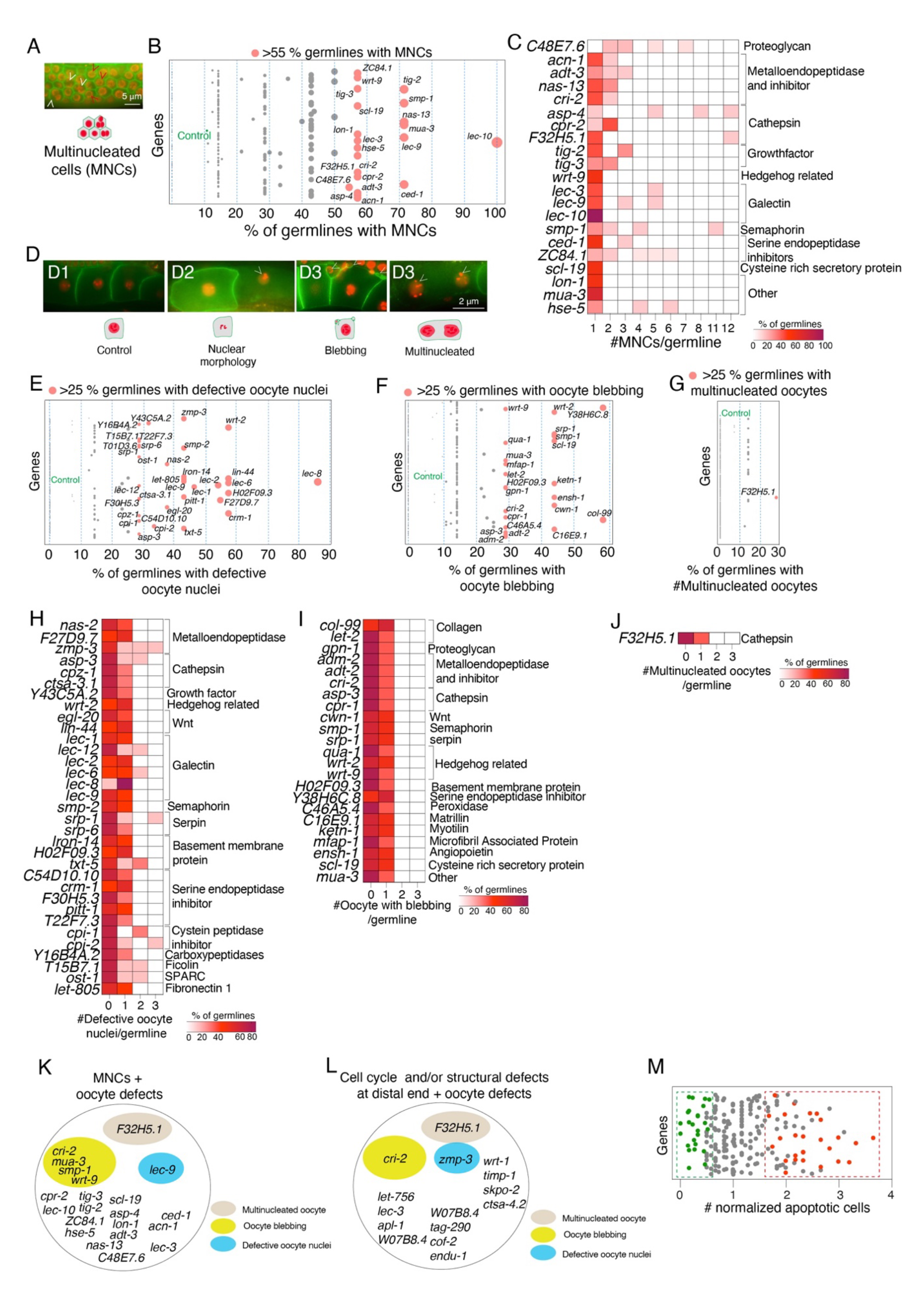
Matrisome control of meiotic germ cell behavior and oocyte health. **(A)** Micrograph showing multinucleated cells (MNCs) in the germline pachytene region. White arrowheads = MNCs. Red arrowheads = normal cells. Scale bar = 5 µm. **(B)** Percentage of germlines with MNCs (circle size proportional to the percentage). Pink circles = significant defect (>55% MNCs compared to controls), grey circles = non-significant. **(C)** Heatmap showing the severity of MNC formation. Each gene knockdown was plotted against the number of MNCs/germline. Color (white to maroon) = % of germ cells with a particular number of MNCs/germline. **(D)** Micrographs showing oocyte defects. D1 = control, D2 = oocytes with defective nuclei, D3 = oocytes with blebbing, D4 = multinucleated oocytes. Defects = white arrowheads. Scale bar = 2 µm **(E-G)** Percentage of germlines with defective oocytes (circle size proportional to the percentage). Pink circles = significant defect (>25% defective oocyte nuclei, oocyte blebbing, or multinucleated oocytes compared to controls), grey circles = non-significant **(H-J)** Heatmaps showing severity of oocyte defects. Gene knockdowns were plotted for defective oocyte nuclei, oocyte blebbing or multinucleated oocytes per germline. Color (white to maroon) = % of germlines with a particular number of oocytes with defects per germline. **(K)** Correlation between MNCs and oocyte defects. 22 gene knockdowns resulted in MNCs, of which 6 had oocyte defects. *F32H5.1* knockdown increased the number of MNCs and multinucleated oocytes. *lec-9* knockdown caused defective oocyte nuclei and MNCs. Silencing *cri-2, mua-3, smp-1,* or *wrt-9* led to an increase in MNCs and oocyte blebbing. **(L)** Correlation between cell cycle and structural defects at the distal end and oocyte defects. From 15 genes required for cell cycle and distal end structure, only *F32H5.1, cri-2* and *zmp-3* showed oocyte defects. **(M)** Graph showing normalized germline apoptosis. Each circle represents the normalized apoptosis after gene silencing compared with control germlines. Green = statistically significant decrease in apoptosis, red = statistically significant increase in apoptosis.

The *C. elegans* germline is self-contained tissue that is capable of cross-talk between distal, proximal and oocyte regions (*7, 12*). We hypothesized that defects observed in distal and pachytene regions would result in unhealthy oocytes. To study oocyte health, we analyzed nuclear morphology, multinucleation, and membrane blebbing in the three oocytes adjacent to the spermatheca (Fig. 3D). First, we determined that defective oocyte nuclei, multinucleated oocytes, and oocyte blebbing were rare defects (∼5%) in control animals (Fig. S10). We classified a gene knockdown significant if it caused >25% penetrance (five times that of control) in oocyte defects (Fig. 3E-G). Using this benchmark, we identified 54 matrisome gene knockdowns that cause oocyte defects, with four genes (*asp-3, wrt-2, srp-1*, and *H02F09.3*) exhibiting multiple oocyte defects (nuclear morphology and blebbing) (Fig. 3E-F). The most common defect was defective nuclei morphology, followed by oocyte blebbing (Fig. 3E-F). Only knockdown of a cathepsin (*F32H5.1*) resulted in multinucleated oocytes (Fig. 3G). Analysis of oocyte defect severity following matrisome gene knockdown showed little difference between each knockdown (Fig. 3H-I). With the exception of *zmp-3* (metalloendopeptidase), all genes showed one (23 genes) or two (9 genes) oocytes with irregular nuclear morphology (Fig. 3H). Additionally, oocyte blebbing and multinucleation were only seen in one oocyte per germline after RNAi (Fig. 3I-J). Taken together, our data suggest that specific matrisome factors control meiosis and oocyte formation, thus extending the role of the matrisome to the proximal germline.

We found that only six out of the 21 genes that showed MNCs resulted in oocyte defects (Fig. 3K). Previous reports suggest that the MNCs are removed by physiological apoptosis in the germline before becoming oocytes (*35*). Except for *F32H5.1*, none of the gene knockdowns causing MNCs produced defective oocytes. In addition, none of the gene knockdowns except for *zmp-3, cri-2 and F32H5.1* that exhibited distal cell cycle and structural defects also showed oocytes defects (Fig. 2C-I, 3L). These data suggest clearance of defective germline cells prior to gamete generation. Indeed, in the hermaphrodite germline ∼50% of germ cells undergo apoptosis to provide cytoplasmic components for maturing oocytes (*36*). We therefore hypothesized that defective germ cells caused by matrisome gene knockdown are cleared by apoptosis before becoming oocytes. To study this, we quantified the normalized number of apoptotic cells per germline compared with the control. We identified 51 gene knockdowns that significantly altered apoptosis (26 upregulated and 25 downregulated), of which 20 were independent of detectable germline defects prior to apoptosis (Fig. 3M, Table S3). While the remaining 31 genes showed apoptosis as well as other defects, strong links between apoptosis and MNCs or cell cycle and structural defects were not visible during the short-term RNAi.

### C-type lectins are essential for *C. elegans* germline function

C-type lectins (CLECs) are calcium-dependent carbohydrate binding proteins that regulate glycosylated proteins (*37*). CLECs contain domains with diverse functions including immune response, apoptosis and cell adhesion (*37*). Based on literature curation and orthology, the *C. elegans* matrisome is predicted to contain > 250 CLECs (*17*). This contrasts the human matrisome with 28 C-type lectins (*38*). Although the presence of a large number of CLECs has not been verified experimentally in *C. elegans*, we systematically silenced each CLEC gene and analyzed the germline. We identified 85 CLEC gene knockdowns (36 in the progenitor zone, 25 in the transition zone, and 24 in both) that showed statistically significant differences in germ cell number at the distal end (Fig. 4A), and 34 gene knockdowns causing MNCs in the pachytene region (Fig. 4B). Silencing of a many CLECs resulted in germlines with several MNCs (Fig. 4C) suggesting a critical function for CLECs in germ cell behavior during meiosis. However, GO coupled with STRING analysis did not reveal any association between CLECs (except *clec-64*) and genes involved in cytokinesis (GO:0000910) and cleavage furrows (GO:0032154, GO:0110070 and GO:0036089). Further, we identified 48 CLEC gene knockdowns leading to oocyte defects (Fig. 4D-F). Interestingly, silencing of 4 CLECs (*clec-12, -185, -240* and *-264*) caused multinucleated oocytes that did not exhibit an increase in MNCs (Fig. 4B and F), suggesting a late-stage defect during oogenesis. While the severity of oocyte defects depended on the CLEC type, only *clec-121, -150, -228*, and -*240* showed multiple oocyte defects (Fig. 4G-I). Finally, we quantified apoptosis following CLEC gene silencing. This analysis revealed that silencing of 43 CLECs resulted in decreased apoptosis (Table S3). This suggests that CLECs promote apoptosis in the germline. Taken together, our data show the that collective functions of CLECs are essential for optimal germline development.

**Fig. 4.**
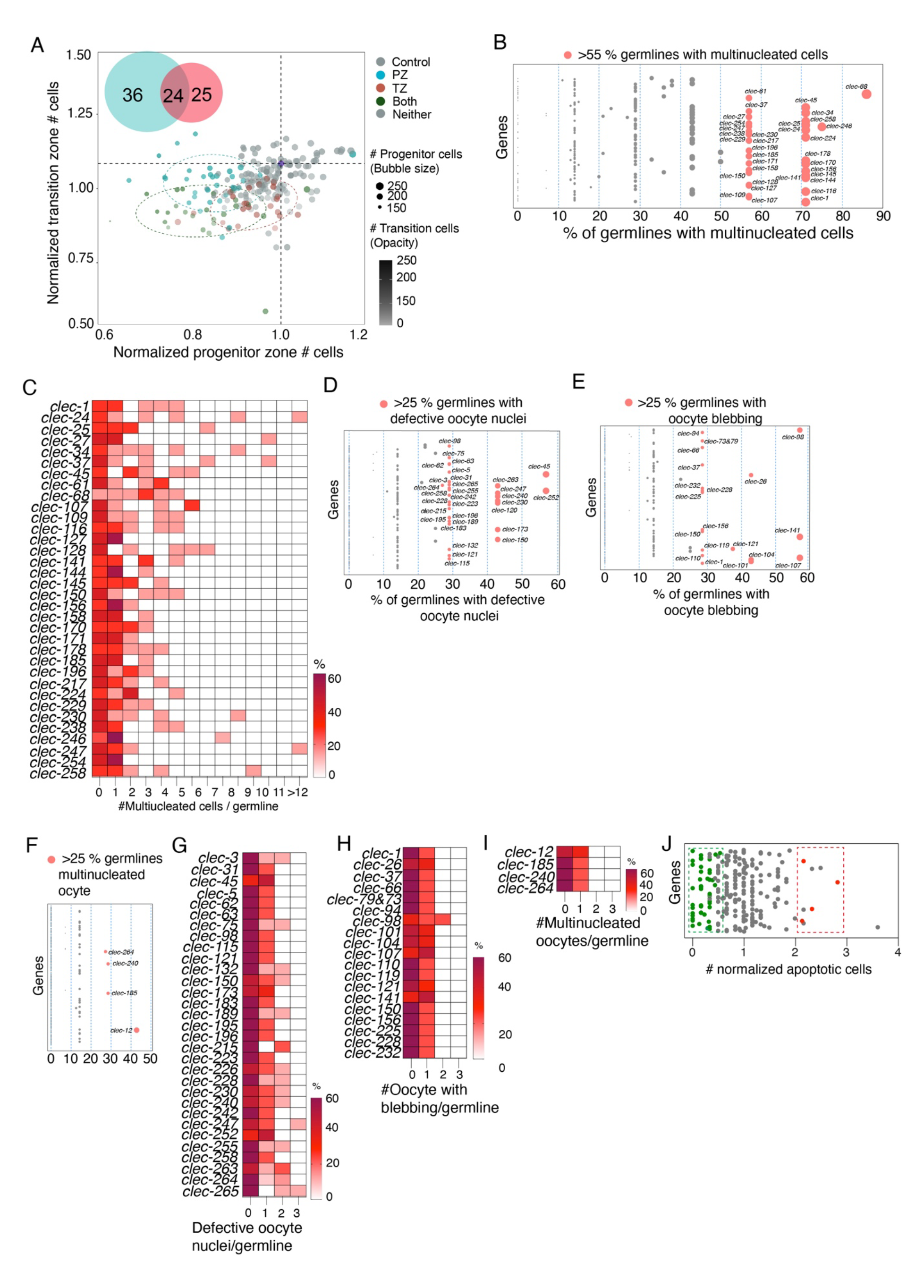
C-type lectin silencing causes diverse germline defects. **(A)** Scatter plot showing germ cell number in progenitor and transition zones after CLEC gene RNAi. Each circle represents a gene. The position of the circle shows the difference in comparison to control RNAi, which is represented as 1. The size and opacity of each circle is proportional to the actual number of germ cells at the progenitor and transition zones, respectively. Knockdown of 36 genes affected PZ cells (blue), 25 affected transitioning cells (red) and 24 affected both regions (green). The size and opacity of each circle indicate the number of germ cells at the progenitor and transition zones, respectively. **(B)** Percentage of germlines with MNCs after CLEC gene knockdown (circle size proportional to the percentage). Pink circle = significant defect (>55% germline MNCs compared to controls), grey circle = non-significant. **(C)** Heatmap showing the severity of MNC formation. Each gene knockdown was plotted against the number of MNCs/germline. Color (white to maroon) = % of germ cells with a particular number of MNCs/germline cells. **(D-F)** Percentage of germlines with defective oocytes (circle size proportional to the percentage). Pink circles = significant defect >25% germline having either defective oocytes nuclei, oocyte blebbing or multinucleated oocytes, grey circles = non-significant. **(G-I)** Heatmaps showing the severity of oocyte defects. Knockdown of each gene was plotted for defective oocyte nuclei, oocyte blebbing, or multinucleated oocytes per germline. Color (white to maroon) = % of germline cells with a particular number of oocytes with each defect per germline. **(J)** Graph showing normalized germline apoptosis. Each circle represents the normalized apoptosis after gene silencing compared with control germlines. Red/Green = Statistically significant increase/decrease in apoptosis, respectively.

### Matrisome interactions and phenotypic mapping reveal a complex functional network

We identified 321 matrisome genes that play important roles in germ cells and/or oocytes (Fig. S11). The germline is a complex tissue maintained by both intrinsic and extrinsic signaling and feedback regulation. To identify the impact of each gene family on the germline, we generated an integrated weighted average plot for each gene family. This showed how each family impacts each germline region considering given the number of genes causing phenotypes and severity of the phenotypes. Thus, a gene family will have high impact if it controls multiple germline regions with high severity, even if the family has a small number of genes. The majority of matrisome genes showed a comparable impact on the germline after RNAi. However, serine proteases, extracellular leucine rich repeats and metalloendopeptidases inhibitors showed higher impact (Fig 5A). To summarize the functions of each gene family at particular germline locations, we determined the number and percentage of genes from each family that generated the phenotype (Fig 5A). As expected, the CLEC family contributed the largest number of genes causing germline phenotypes, followed by metalloendopeoptidases, serine endopeptidases inhibitors, cathepsins and galectins (Fig 5A). The percentage revealed the proportion of genes (regardless of the total number in the family) from each family that resulted in a phenotype at a particular germline region (Fig 5A).

**Fig. 5.**
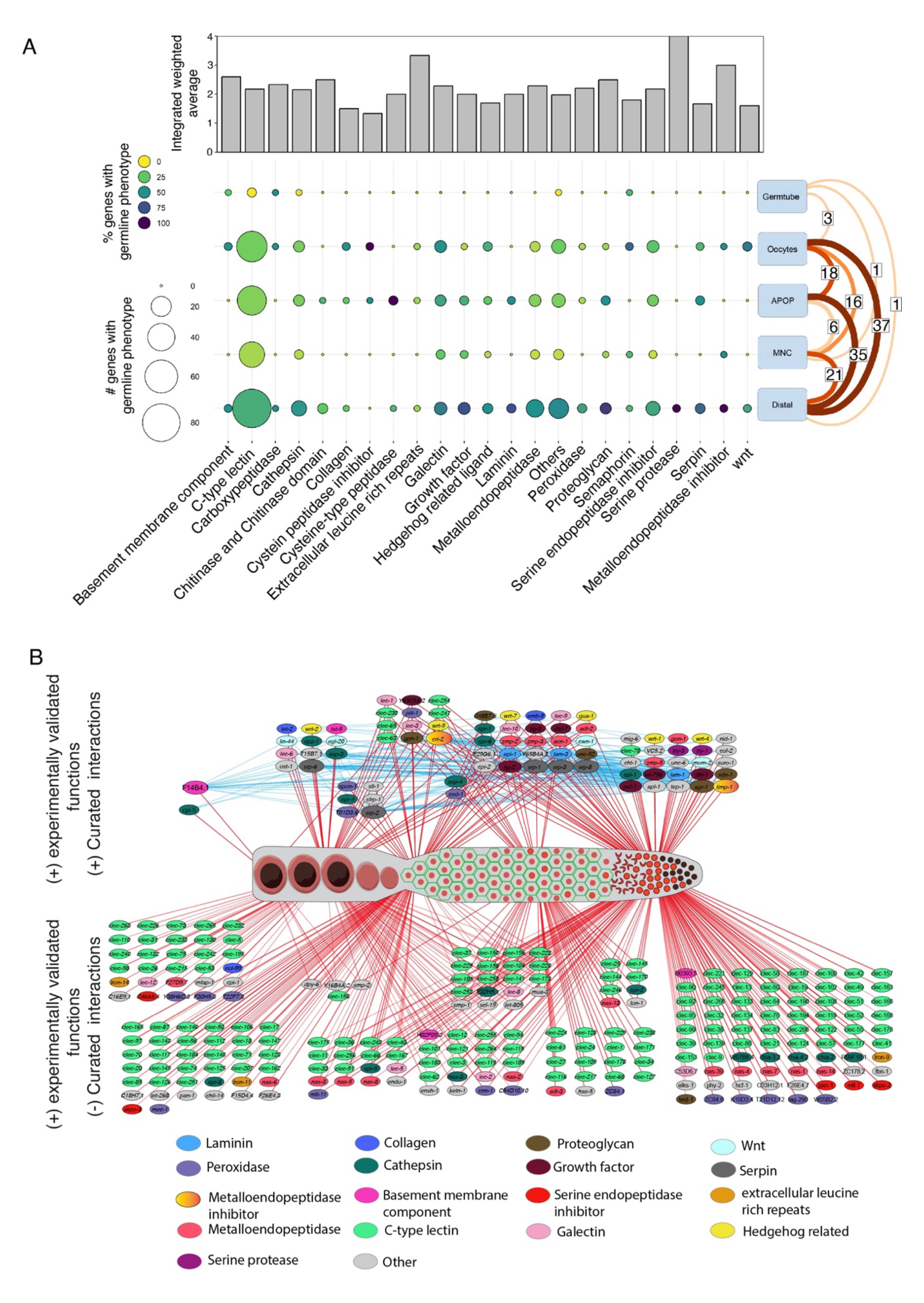
Mapping matrisome interactions and associated germline functions. **(A)** Top panel – Bar graph showing the impact of each gene family in the germline, calculated by weighting the phenotype average based on the number of genes causing the phenotype. Left axis – Integrated weighted average. 0 to 4 = low to high impact. Bottom panel - balloon plot showing the distribution of genes causing phenotypes in each germline region. Size of the circle = number of genes showing a phenotype. Color of the circle = percentage of genes in each family showing a phenotype. The numbers on the right of the plot shows the number of genes that cause phenotypes in multiple regions. **(B)** Map showing the curated interactions between matrisome genes with experimentally verified phenotype. Top schematic - network showing all the genes regulating the germline and have curated interactors. Blue lines show interactions between genes, and the red lines show the location of germline functional impact. Size of the node = proportional to number of interactions. Each gene family represented by node color. Gene families with less than two members indicated as other. Bottom schematic – Map of matrisome RNAi phenotypes, mainly constituting CLECs, that did not reveal interactions with other matrisome genes in validated databases.

Matrix digestive enzymes, such as metalloendopeptidases, cathepsins, and serine proteases, also had profound effects on the germline (Fig. S4), suggesting a critical role of matrix remodeling in germ cell maintenance. The important core matrisome genes (collagen, laminin, nidogen, and proteoglycans), many of which are either substrates or regulators of matrix digestive enzymes, also showed germline defects (*39, 40*). Moreover, interactions between these genes may determine the matrisome functional output. We therefore examined how these putative interactions and phenotypes are linked. We performed STRING analysis of matrisome genes we found are important for germline development, with curated interactions and mapped them according to the experimentally validated phenotypes (Fig. 5). Curated interactions were generated only from validated experiments and functional databases with associated references.

This analysis identified 76 matrisome genes with validated germline phenotypes that could be placed in an interaction network formed of 305 unique interactions (Fig. 5 and S12). Further, we discovered 21 (of 76) matrisome factors that have >10 other matrisome interactions that also are required for germline development (Fig. S12). Two inhibitors of metalloproteinases, *timp-1* and *cri-2*, showed the highest number of interactions with other matrisome genes with validated phenotypes, followed by the basement membrane component *F14B4.1* (Fig. S12). These interactions were generated based on curated data and do not guarantee phenotypic links. To study the association between the interactions and phenotypes, we compared the experimentally generated phenotypes of each gene with those of their interacting partners. We identified 60 out of 76 interacting genes (79%) that had at least one shared germline phenotype with their interacting partners following gene silencing (Fig. S12). These 60 genes represented 157 of total 305 interactions between matrisome genes (51%) (Fig. S12). Silencing the metalloproteinase inhibitors *cri-2* and *timp-1* resulted in four and two germline defects, respectively (Fig. S13). The majority of their interacting partners, including their substrates (metalloendopeptidases), showed same defects upon knockdown (Fig. S13) (*41*). The basement membrane component *F14B4.1*, showed the second highest number of interactions. However, the only phenotype resulting from silencing *F14B4.1* was defects in germ tube morphology (Fig. S14), which was not shared by any of the interactors (Fig. S14). Silencing the matrix structural protein laminin (*epi-1, lam-1* and *lam-3*) caused defects at the distal end and an increase in apoptosis (Fig. S15). These defects are shared by many of its major interactors, including proteoglycans. Proteoglycans are a group of receptors present both on the cell surface and in the extracellular matrix, where they act as receptors for a plethora of other matrisome molecules (*39*). Our analysis verified that the resulting phenotypes after RNAi were similar between the proteoglycans (*agr-1, gpn-1, sdn-1* and *unc-52*) and their interactors (Fig. S15). Another group of proteoglycan ligands are growth factors, some of which are known to regulate germline signaling (*42*). We found that the growth factors (*daf-7, dbl-1, let-756, tig-2* and *tig-3*) and their interactors, all control germ cell number, MNCs and apoptosis (Fig. S15). We found that multiple serpins, a CD109 molecule (*tep-1*), and amyloid β-precursor like proteins (*apl-1*) also have large number of interaction partners among the matrisome genes. Interestingly, curated interactions between CLECs and other matrisome molecules are minimal, even though they are the largest family of matrisome molecules. Similarly, we identified widespread roles for galectins in the germline, although their interactions with the other matrisome genes important for germline development were limited. Finally, we analyzed the functional pathways controlled by the genes in the network using the Reactome database (*43*). Overrepresentation analysis of comparable mammalian orthologs suggest that 44 unique matrisome genes contribute to 170 signaling pathways. Among these, 15 matrisome genes were found on four pathways with a high confidence level (Fig. S17). Overall, we identified a subset of genes that have physical and/or functional interactions in curated databases that produce specific local or global germline phenotypes after gene silencing. This suggests that an intricate network of molecules forms a matrisome landscape to control germ cell fate.

## Discussion

Our systematic silencing of matrisome genes in *C. elegans* provides insight into how the matrisome regulates germ cell fate *in vivo*. Our approach of short-term gene silencing allowed the acute requirement of matrisome factors to be assessed, without long-term secondary impacts on development. In addition, by employing the *C. elegans* germline model, we could study cell fate in self-contained tissue (*12, 30*). While cells are contained in the germ tube, they receive signals from other tissues and extracellular structures (*6*). The comprehensive analysis of phenotypes resulting from silencing matrisome genes reveals a complex functional network of molecules that control germ cell proliferation, differentiation, and apoptosis, to produce healthy oocytes (*30*). We found that changes in matrisome homeostasis can affect one or more of these germline processes. Cells in the progenitor zone undergo mitotic cell division to maintain the germ cell population (*44*). However, understanding how the matrisome controls the mitotic cell cycle is limited. Here, we revealed association between matrisome gene function and the cell cycle of mitotic germ cells, which likely depend on the structure of the distal germline. This provides an additional layer of signaling in the *C. elegans* progenitor zone, which has not been explored previously. Similarly, we identified roles of matrisome genes in the pachytene region, where loss of matrisome gene function causes inappropriate multinucleation of meiotic cells, indicating that some matrisome genes have roles in cytokinesis and/or cleavage furrow formation. Organisms possess mechanisms such as apoptosis that protect their germline from inheritance of defective gametes. Therefore, termination of germ cells with defects such as multinucleation is key to maintaining the fidelity of the next generation (*35, 45*). Our attempt to link early germ cell defects to later apoptosis was not successful for two main reasons. First, a population of gene knockdowns showed oocyte defects, independent of early germline defects. Second, silencing of some genes caused changes in apoptosis independent of other defects. It is likely short time of gene silencing did not allow other mechanisms take into effect.

A reduction in the number of germ cells or defective oocytes should result in reduced fertility. Previous studies showed that loss of certain matrisome genes results in reproductive disorders in *C. elegans* and other organisms (*7, 40, 46*). Our study provides the likely cellular causes underpinning these observations. Our collective data reveal the complex functions of matrisome factors. With a limited number of interactions in curated databases, we attempted to map these functions into an interaction network. Admittedly, a number of matrisome factors with experimentally validated functions require further interaction studies, which are beyond the scope of this research. Our data show that silencing genes that encode extracellular matrix remodelers had the greatest impact on germ cell development. This includes proteolytic enzymes (e. g. metalloendopeptidases and cathepsins) and their regulators that can remodel structural proteins such as collagens and laminins, after their deposition into the extracellular space (*47*). Additionally, extracellular matrix modifications can alter ligand presentation to cells (*48*). This suggests that matrisome modifications are continually required for controlling cellular fates in tissues. However, we found that silencing of structural molecules such as laminins and collagens resulted in weak defects in the germline. This is likely a function of the short-term silencing and abundance of these proteins that are pre-deposited during development and stable during the window of analysis owing to their slow turnover (*49*). We found that growth factors and their likely receptors (proteoglycans) mostly function in the distal pachytene region of the germline, aligning with their previously ascribed roles (*6, 7*). Similarly, we confirmed the previously identified role of hedgehog related genes in the distal germline while identifying additional roles at the proximal and oocyte regions (*50*). We also found widespread novel roles for lectins (galectins and CLECs) in the germline. While the number of predicted CLECs differs between human and *C. elegans* matrisomes, galectin composition is comparable (*17, 38*). Nevertheless, silencing of these genes produces a wide variety of defects in the germline. Both galectins and CLECs can bind to carbohydrates, thereby regulating glycosylated proteins (*37, 51*). Many animal proteins are glycosylated, and thus galectins and lectins may broadly regulate their function. This could be why loss of multiple lectins affected germ cell behavior. However, we were unable to show this on the interaction-phenotype map because of the lack of verified studies. Taken together, our data reveal an exquisite requirement for a balance in the matrisome landscape to control germ cell fate and maintenance in the *C. elegans* germline. Importantly, the lack of germline expression for many matrisome genes suggests that they regulate germ cell fate non-cell-autonomously. Thus, the effect of environmental, stress and aging signals on germ cell behavior are likely coordinated by controlling matrisome expression and release from distal tissues (*52*). Finally, given that our analysis of matrisome gene function was after the early developmental stages were complete, our data reveal a continual requirement for remodeling of the matrisome landscape for faithful germ cell behavior and gamete generation.

## Materials and Methods

### *Caenorhabditis elegans* breeding

*C. elegans* strains were maintained on Nematode Growth Medium (NGM) plates and fed with OP50 *Escherichia coli* bacteria at 20°C, unless otherwise stated.

### Plasmid production for RNAi

RNAi plasmids for 333 genes were available commercially. The plasmids were verified by Sanger sequencing. 110 plasmids were cloned in house. The sequence for each gene was amplified using forward and reverse primers. HindIII sites were introduced at both ends while amplification of the gene sequence. The empty vector (L4440) and amplified product were digested using HindIII enzyme. The amplified gene was ligated into the vector using DNA ligase enzyme. The plasmids with verified sequences were transformed into HT115(DE3) *E. coli* bacteria for RNAi experiments.

### RNA interference experiments

HT115(DE3) E. coli bacteria expressing RNAi plasmids for specific genes or an empty vector (L4440) were grown in Luria Broth (LB) + Ampicillin (100μg/ml) at 37°C for 16hr. Saturated cultures of RNAi bacteria were plated on RNAi plates and allowed to dry for at least 24hr. L4 hermaphrodites were placed on the RNAi plates and incubated for 16hr at 20°C before proceeding with germline analysis.

### Distal cell number analysis

Cell number at the progenitor and transition zones were analyzed by DAPI staining and 3D modelling. Germlines were extruded from sedated worms and fixed on a poly-L-lysine coated slide using ice cold methanol for 30 seconds and then in 4% paraformaldehyde (PFA) for 30min. Fixed germlines were washed twice in phosphate buffered saline (PBS, pH 7.4) containing 1% Tween 20 (PBST) and blocked using 30% normal goat serum. The germlines were incubated with 4′,6-diamidino-2-phenylindole (DAPI) for 2hrs at room temperature. After staining, germlines were washed twice with PBST. Slides were mounted by applying a drop of Fluroshield mounting media (Sigma) on the germlines, followed by a coverslip. The germlines were imaged using confocal microscopes (Leica SP8 microscope at 63x objective, Olympus A1RHD microscope at 40x objective or Leica Stellaris microscope at 40x objective). The images were converted into 3D models by using Imaris 10.0 software as described previously *(7, 21)*. Briefly, the diameter of DAPI stained nuclei in the mitotic region were defined with an XY diameter, while transition zone nuclei had an XY and Z diameter. The diameters were decided according to the objective used. The accuracy of diameter was confirmed using control RNAi samples so that the previously observed values as in wild-type germline were obtained *(7)*. The bubble plot showing progenitor and transition zone cell number was created using the ggplot2 of the programming language ‘R’ *(53)*. The cell numbers were plotted showing progenitor zone cell number on X-axis and transition zone cell number on Y-axis. The cell numbers were normalized against control RNAi values, which is 1. Actual values are represented as by the size (progenitor zone) and opacity (transition zone) of each bubble on the graph.

### Cell cycle analysis

Cell cycle analysis was performed by 5-ethynyl-2′-deoxyuridine (Edu) labeling via soaking. After RNAi feeding, worms were washed in a solution of M9 containing 0.01% Tween-20. An equal volume of EdU and M9 buffer were added to the wash solution to create a final EdU concentration of 250μM. Worms were left in solution and rotated for 15 min before being placed on an unseeded agar plate and left to recover for 1-2min. Germlines were then extracted and fixed as described above. EdU-labelled cells were stained with a Click-iT Alexa Flour (488) EdU labelling kit (Invitrogen) by undergoing 2x 30min Click-iT reactions followed by 2hr DAPI staining. After staining, germlines were washed twice with PBST. Slides were mounted by applying a drop of Fluroshield mounting media (Sigma) on the germlines, followed by a coverslip. Edu labelled germlines were imaged using Leica SP8 at 63x or Olympus A1RHD microscopes at 40x objective and analyzed using Imaris 10.0 software. The stages of cell cycle were identified based on intensity and shape of Edu staining (Fig. 1) *(26)*. By using Imaris, all the cells stained with Edu were marked automatically. The cells were manually observed to ensure the correctness of Imaris-based identification. To avoid the error in manual quantification of Edu staining intensity and variations between runs, intensity was calculated using Imaris Software. The cells with highest intensity (top 30% of maximum intensity threshold) were identified as mid and late S phase (Fig. 1) *(26)*. The two stages were distinguished manually by counting the cells in the late S-phase. Late S phase shows a punctate staining in the nuclei, which were counted and subtracted to obtain mid S phase cell number. The cells with low intensity (bottom 70% or less) were identified as early S-phase. The values were normalized against the values from control RNAi values.

### Cytoskeleton analysis

For cytoskeletal staining, germlines were extruded and fixed in 4% paraformaldehyde (PFA) for 30min. The germlines were then washed in PBST and incubated with phalloidin (cytoskeletal) and DAPI (nuclei) for 2 hr before washing with PBST. The germlines were then imaged as explained above. The phenotypes were analyzed by observing the cytoskeleton structure at the distal end using Imaris 10.0 software. Gene knockdown was considered significant if the percentage of germlines showing cytoskeletal defects was 50% or more after RNAi.

### GLP-1 and nuclei distribution analysis

Germlines were extruded from the *glp-1::v5(q1000)* strain after RNAi. This strain expresses GLP-1 with a V5-tag *(29)*. The isolated germlines were fixed and blocked as explained above. The slides were washed twice in PBST and incubated with 30% normal goat serum containing diluted antibody against V5-tag overnight at 4°C. After incubation, germlines were washed twice with PBST and incubated with fluorophore-conjugated secondary antibody and DAPI for 2hrs. The slides were then washed, mounted and imaged, and 3D models of DAPI staining were generated using Imaris 10.0, as described. To create a 3D model of GLP-1, the surface function of Imaris 10.0 was used. For each knockdown, respective control RNAi germlines were used to define surface development parameters. The background, minimum and maximum intensity threshold, detailing of surface were defined for germline after control RNAi. To avoid the variations in staining quality and intensity, each parameter was defined for a particular run using control samples and kept uniform during the 3D model creation of all samples. Gene knockdown was considered significant if the percentage of germlines showing variations in GLP-1 phenotype was 50% or more after RNAi. To study the distribution of nuclei, the distance between each nucleus and its closest neighbors were analyzed using Imaris distance function. First, using the 3D model of the nuclei, the automated analysis calculated the average of three distances for every nucleus. The mean was used to define the distribution of nuclei in the germline. A smaller distance indicates closer packaging of nuclei at the progenitor zone. To measure the distance between each nucleus and GLP-1, the distance from the center of the nucleus to the nearest GLP-1 surface was measured using Imaris. The average of the distances between all nuclei and their closest GLP-1 surface was used to define the proximity of GLP-1 to the nucleus at the progenitor zone.

### Analysis of MNCs, apoptosis, oocyte defects and germtube defects

For analyzing the proximal phenotypes, *unc-119(ed3) III; ltIs37 IV; ltIs38* strain was used. This strain expresses mCherry tagged Histone 2B (nucleus) and GFP fusion binding PI4, 5P_2_ (plasma membrane), which allows observation of nuclei and the plasma membrane in live animals *(30)*. Germlines were imaged using Zeiss Axiocam microscope at 63x at different Z planes. Each germline was manually analyzed for phenotypes (iii to viii) listed in Fig S3 *(30)*. The percentage of germlines with MNCs and oocyte defects was calculated. If a gene knockdown showed defects 5 times more than control, it was identified as significant. To investigate the severity of MNCs and oocyte defects, the number of defects per germline was calculated. Apoptosis was quantified by counting the nuclei with morphology as explained previously *(30)*. Briefly, the apoptotic cells were recognized by a condensed nuclei morphology and strong mCherry (Histone 2B) intensity in *unc-119(ed3) III; ltIs37 IV; ltIs38* strains. They also show irregular cell boarders under differential interference contrast (DIC) microscopy. Apoptotic cells were first identified by mCherry level and then confirmed with DIC microscopy. Normalized apoptosis after knockdown was calculated by using values obtained from control RNAi from 302 germlines. The germtube defects were calculated by manually counting the germline that do not maintain the U shape of germtube in the worm (phenotype viii of Fig S3)

### Gene Ontology analysis

Gene ontology search was performed using QuickGo - A web-based tool for Gene Ontology searching *(54)*. The following terms were used in this study -Germ cell development (GO:0007281), gamete generation (GO:0007276), mitotic cell cycle (GO:0000278), cytoskeleton (GO:0015629), cytokinesis (GO:0000910), cleavage furrow (GO:0032154), cleavage furrow formation (GO:0036089), cellularization of cleavage furrow (GO:0110070).

### Generation of heatmaps and balloon plots

The heatmaps for figure 1F were constructed using Complex Heatmap package of R language. Other heatmaps were generated using GraphPad Prism 7.0. The balloon plot was visualized using ggplot2 package of R language *(55)*

### Generation of STRING network

The STRING network was constructed based information fetched from STRING database *(56)*. The network only included the experimentally verified and curated interactions. For Fig. 2B, the network was constructed from a combined list of two gene sets. One set of genes were associated with mitotic cell cycle and the second set of genes with experimentally verified distal phenotypes. Similarly, Fig. S7B was constructed from genes associated cytoskeleton and with experimentally validated progenitor zone. Finally, for Fig. 5B was constructed with experimentally verified matrisome gene list and the network was visualized using Cytoscape software (version 3.10.0) *(57)*.

### Integrated weighted average calculations

The integrated weighted average was calculated for each family of matrisome genes according to the following:

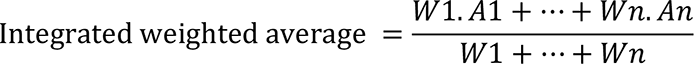

A= average phenotype

W= weight of the genes of the family in each region calculated as follow:

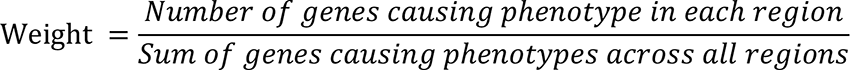

### Quantification and statistical analysis

Statistical analyses were performed in GraphPad Prism 7 using one-way analysis of variance (ANOVA) for comparison followed by a Dunnett’s Multiple Comparison Test where applicable. Unpaired t-tests were performed if the comparison was for two conditions. Values are expressed as mean ± S.E. Differences with a P value <0.05 were considered significant.

## Acknowledgments

We thank Prof. John Couchman, Prof. Anders Malmstrom and Prof. Gunilla Westergren-Thorsson for advice and comments on the manuscript. We thank Lisa Karlsson for generating schematics. Imaging for this project was performed at Monash Microimaging, Lund Bioimaging Center and Lund Stem Cell Centre Imaging Facility. Some strains were provided by the *Caenorhabditis* Genetics Center (University of Minnesota), which is funded by the NIH Office of Research Infrastructure Programs (P40 OD010440). The gene family classification was performed as curated in the Wormbase.

## Author contributions

Conceptualization: SG, RP. Methodology: AA, LP, JF, RG, SG. Investigation: AA, LP, JF, SG. Visualization: AA, RG, SG. Funding acquisition: SG, RP. Project administration: SG. Supervision: SG, RP. Writing – original draft: SG. Writing – review & editing: SG, RP, AA, LP, JF

## Data and materials availability

All data is available in the main text or supplementary materials. In addition, Source Data are provided with this paper. There are no accession codes, unique identifiers, or weblinks in our study and no restrictions on data availability. Materials will be available upon request from the Gopal laboratory.

