## Supplemental figures and tables for "A matrisome atlas of germ cell development"

[2] Development and Stem Cells Program, Monash Biomedicine Discovery Institute. Department of Anatomy and Developmental Biology, Monash University, Australia.

* Correspondence should be addressed to S.G. and R.P.
E mail:

† Equally contributed

### Supplemental data

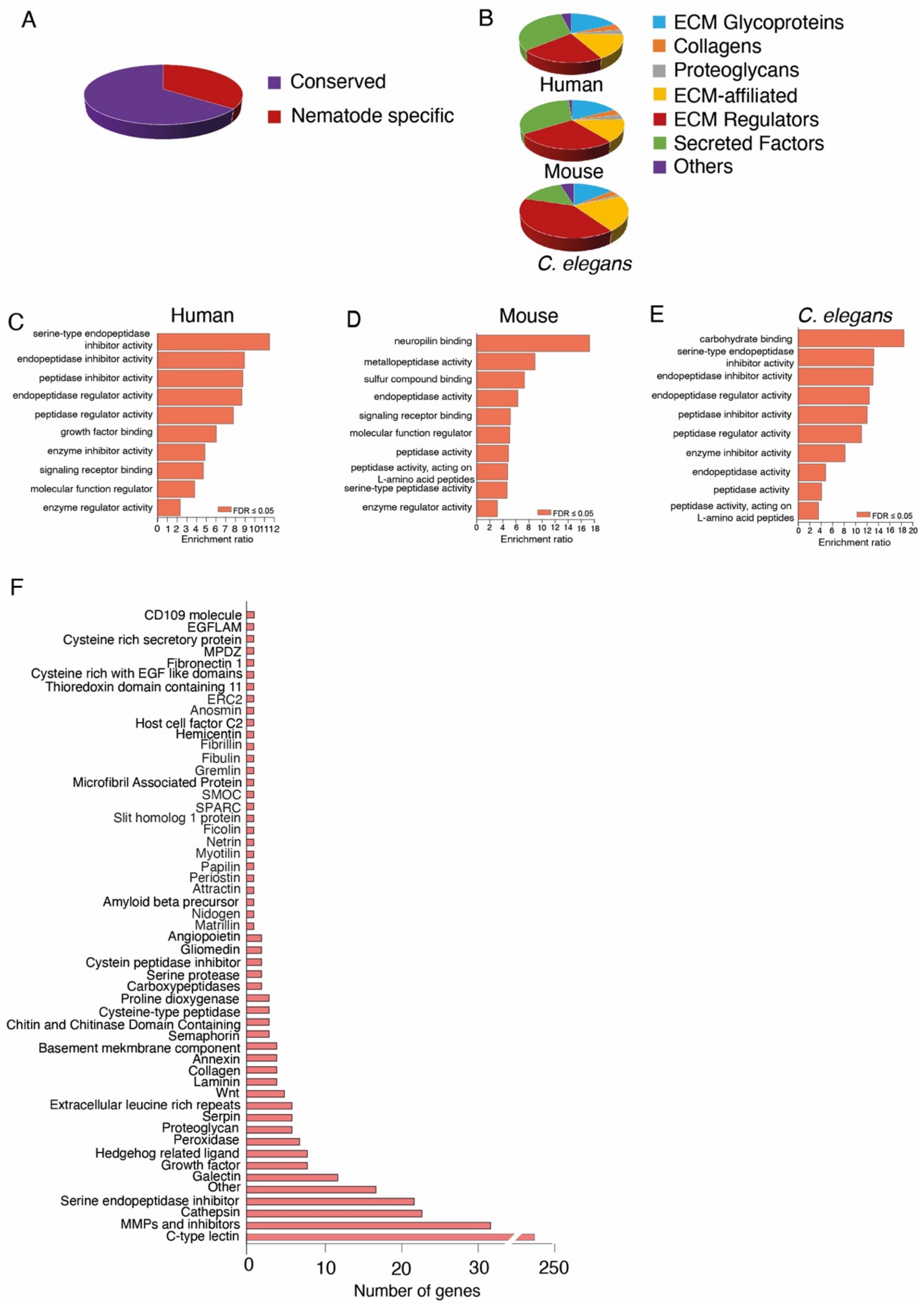


**Fig. S1. Conservation of the *C. elegans* matrisome.** **(A)** Number of conserved and non-conserved matrisome genes in *C. elegans*. **(B)** Composition of the major protein families within the matrisome of humans, mice and *C. elegans*. **(C-E)** Gene Ontology analysis of humans, mice, and *C. elegans* matrisome proteins showing similar biological processes controlled by the matrisome. **(F)** The gene families constituting the conserved *C. elegans* matrisomes according to Wormbase data.

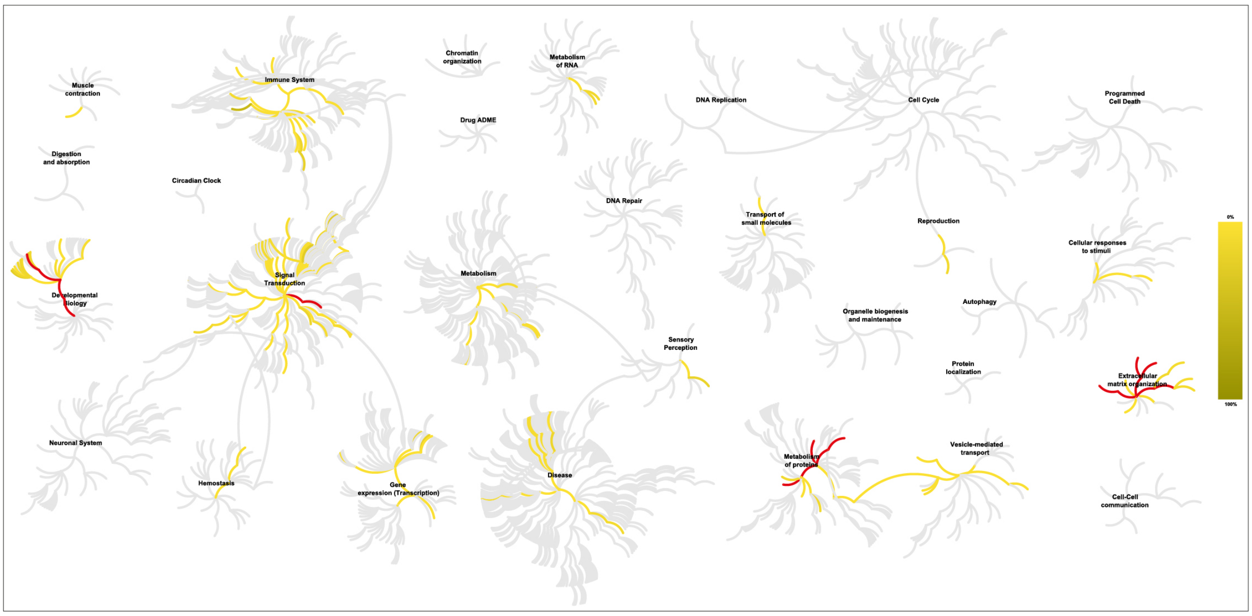


**Fig. S2. Functional diversity of the *C. elegans* matrisome.** Overrepresentation analysis of human orthologs of conserved *C. elegans* matrisome using the Reactome database showing their roles in 250 signaling pathways. Fourteen pathways with high confidence ($P < 0.05$) are shown in red.

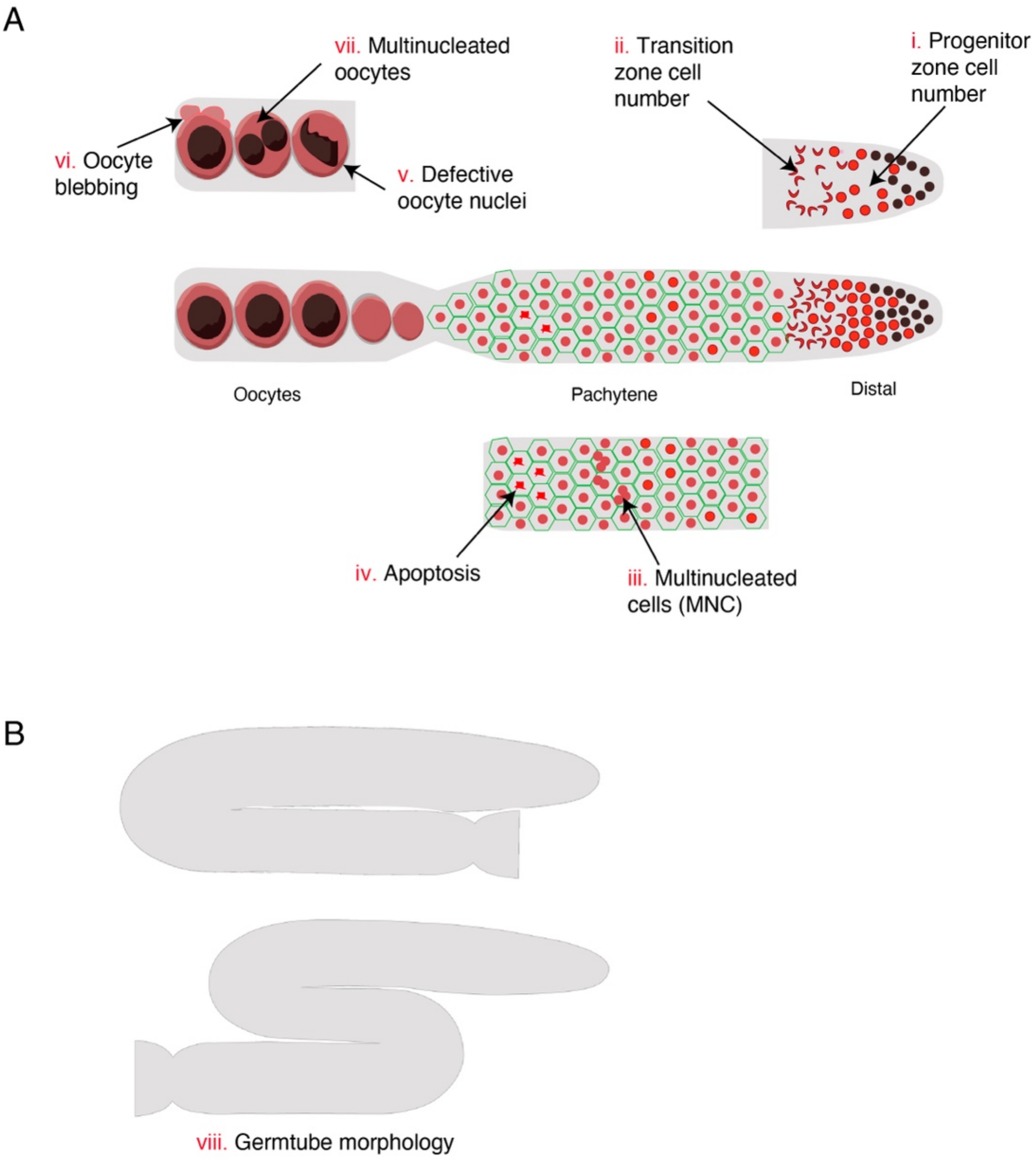


**Fig. S3. Germline phenotypes analyzed. (A)** Germ cells and oocytes were analyzed for 7 distinct defects. i - Progenitor zone cell number, ii - transition zone cell number, iii - multinucleated cells at pachytene, iv - apoptosis at late pachytene, v - defective oocyte morphology, vi - oocyte blebbing, vii - multinucleated oocytes. **(B)** viii - germ tube morphology. Normal germline organization in a U-shape (top). Example defective germline organization following RNAi (bottom).

| Gamete generation GO:0007276 | | | | |
|---|---|---|---|---|
| ***C. elegans*** | **Human** | **Mouse** | **Zebrafish** | **Drosophila** |
| *cwn-1* | WNT4 | Wnt4 | wnt4 | |
| *dbl-1* | BMP4 | Bmp4 | | dpp |
| *tig-2* | BMP8A | Bmp8a | | gbb |
| *let-268* | | | | Plod |
| *pvf-1* | | | | Pvf1 |
| *adt-3* | | | | stl |
| *nas-39* | | | | tld |
| *cri-2* | | | | Timp |
| *skpo-2* | | | | Pxt |
| *mlt-7* | | | | Pxt |
| *cpl-1* | CTSL | Ctsl | | |
| *gon-1* | ADAMTS1 | Adamts1 | adamts1 | AdamTS-A |
| *acn-1* | ACE | Ace | ace | |
| *zmp-3* | MMP19 | | | |
| *zmp-5* | MMP19 | | | |

| Germ cell development GO:0007281 | | | | |
|---|---|---|---|---|
| ***C. elegans*** | **Human** | **Mouse** | **Zebrafish** | **Drosophila** |
| *cwn-1* | WNT4 | Wnt4 | wnt4 | |
| *dbl-1* | BMP4 | Bmp4 | | dpp |
| *tig-2* | BMP8A | Bmp8a | | gbb |
| *let-268* | | | | Plod |
| *pvf-1* | | | | Pvf1 |
| *adt-3* | | | | stl |
| *nas-39* | | | | tld |
| *cri-2* | | | | Timp |
| *skpo-2* | | | | Pxt |
| *mlt-7* | | | | Pxt |

**Fig. S4. Association of matrisome genes with gamete generation and germ cell development.** 15 matrisome genes with experimentally validated phenotypes were present in genes annotated with gene ontology terms ‘gamete generation’ (15 genes) and ‘germ cell development’ (10 genes) in *C. elegans*, human, mouse, zebrafish, and *Drosophila*.

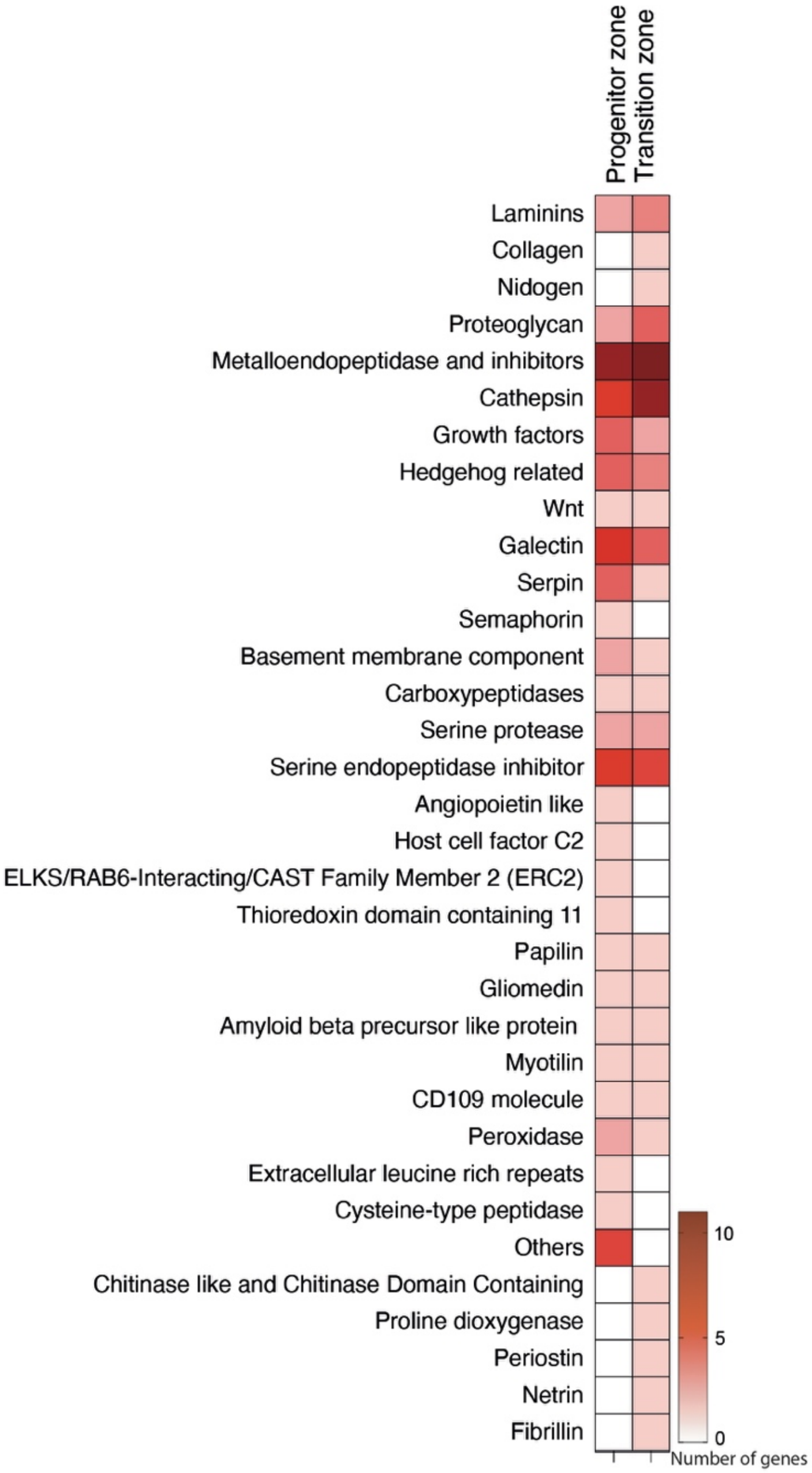


**Fig. S5. Heat map depicting gene families functioning in the distal germline.** The number of genes from each matrisome family that produced a change in germ cell number in the progenitor and/or transition zone is shown.

A

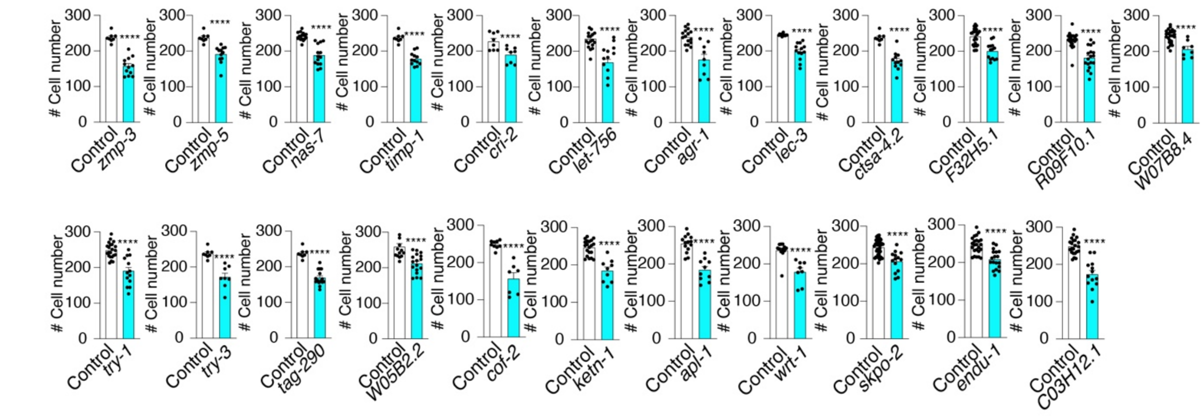


B

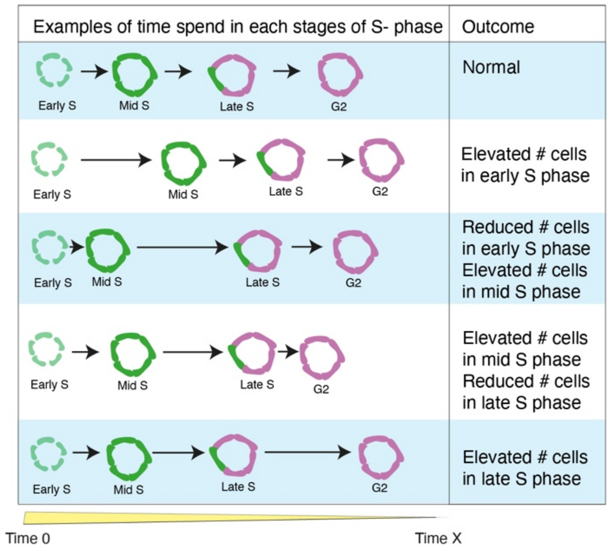


C

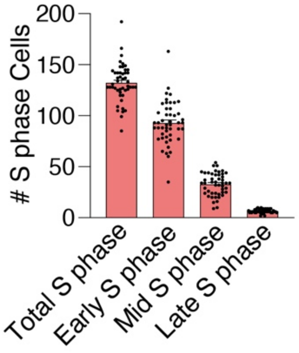


D

| | |
|---|---|
| *apl-1*<br>*let-756*<br>*kten-1* | Known genes associated with mitotic cell cycle |
| *cri-2*<br>*W07B8.4*<br>*wrt-1*<br>*timp-1*<br>*F32H5.1*<br>*skpo-2*<br>*endu-1*<br>*lec-3*<br>*zmp-5*<br>*zmp-3* | Novel genes associated with mitotic cell cycle at the progenitor zone |

**Fig. S6. Cell cycle analysis in the progenitor zone. (A) Graphs showing the genes with the most significant reduction in the number of germ cells in the progenitor zone. ****P>0.0001. (B) Examples of EdU staining and quantification of the S phase stages.** Time each cell spends in different stages of the S phase determines the total number of cells in a particular stage. **(C)** Graph showing the total number of cells in S phase and cells at specific stages of S phase calculated based on EdU staining in the control (n >120). **(D)** Genes associated with the mitotic cell cycle control. Previously reported genes are shown in blue and novel genes are shown in grey.

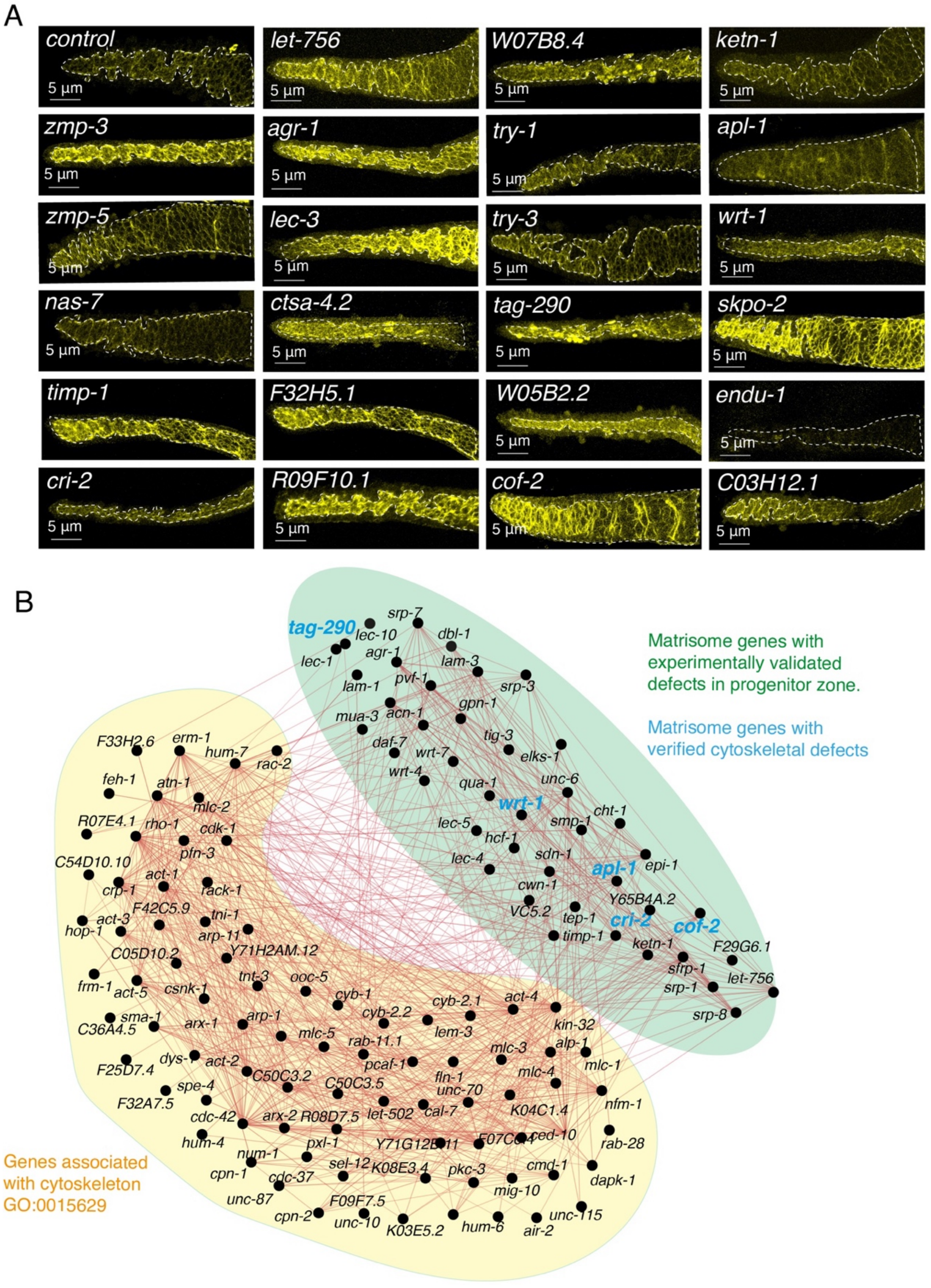


**Fig. S7. Matrisome association with cytoskeleton structure. (A)** Phalloidin staining showing the cytoskeletal structure at the distal end of control and matrisome gene-silenced germlines. **(B)** Interaction analysis of the combined list of genes that showed a phenotype at the distal end and genes annotated with the gene ontology term ‘cytoskeleton’ (GO:00115629). Green shade = genes with experimentally verified changes in germ cell number at the distal end. Yellow shade = genes from the gene ontology list GO:00115629 (cytoskeleton). Experimentally confirmed genes with roles in the germline cytoskeleton are shown in blue.

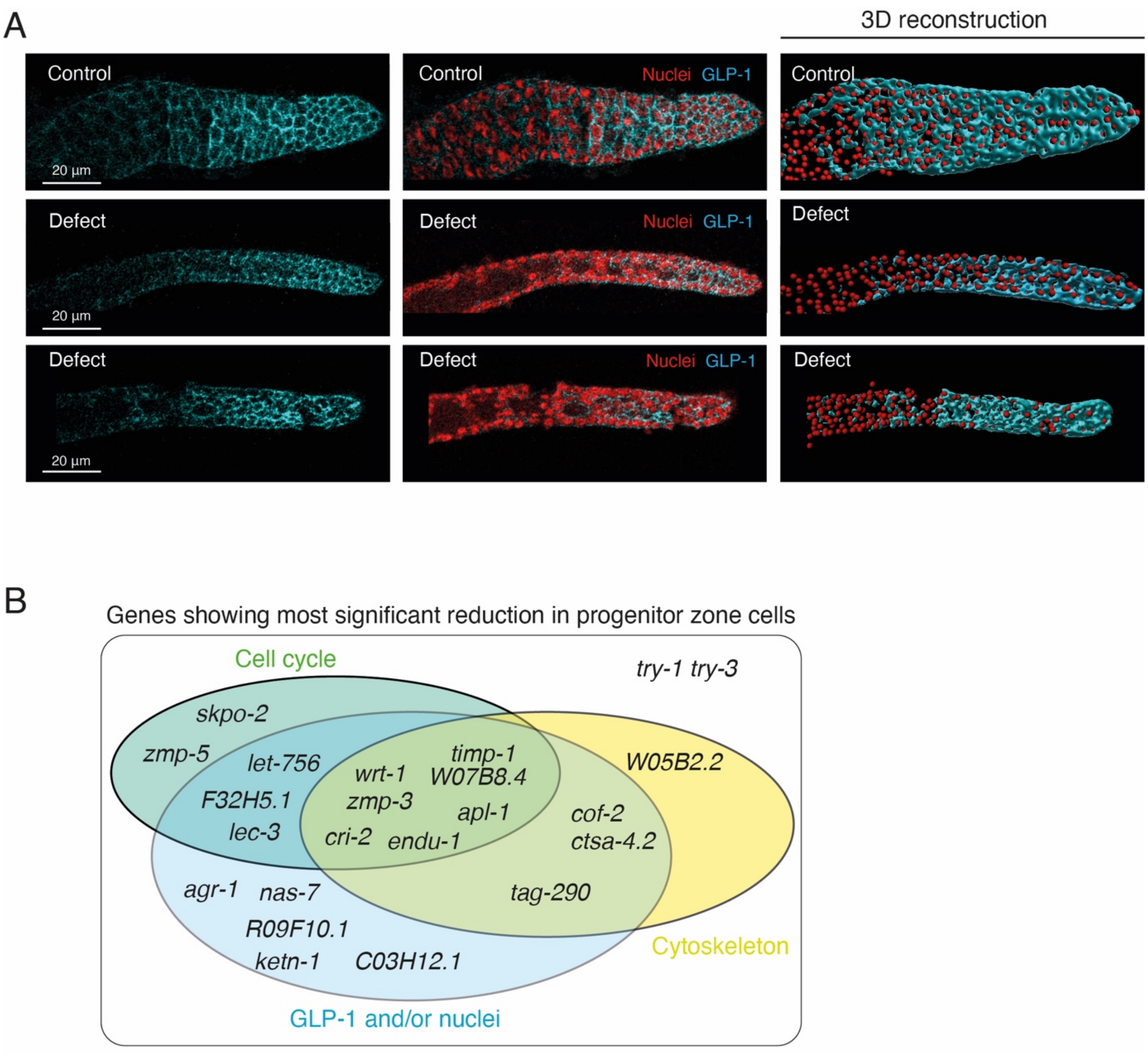


**Fig. S8. GLP-1 expression and phenotypes in the progenitor zone. (A)** Representative images showing control and defective germlines stained for GLP-1 protein (blue) and DAPI (red). Top panel = control germline. Middle and bottom panels = defective germline after RNAi. 3D reconstruction shows GLP-1 localization with respect to nuclei. (**B**) Ven diagram showing the impact of most significant genes on cell cycle, cytoskeleton and GLP-1/nuclei in the progenitor zone.

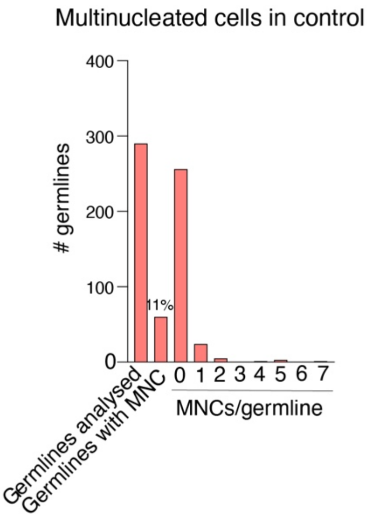


**Fig. S9. Analysis of multinucleated cells (MNCs) in control germlines.** Graph showing the total number of germlines (n = 290) analyzed and germlines with MNCs. 11% of germlines with showed MNCs after control RNAi. The number of MNCs per germline ranged between 1 and 7

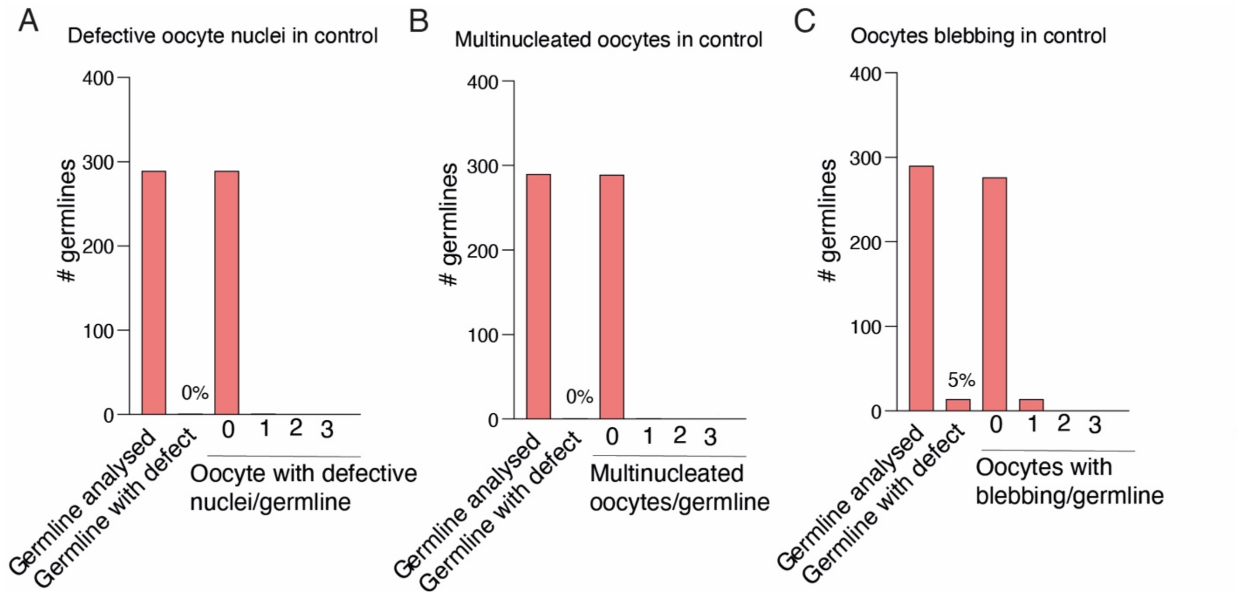


**Fig. S10. Analysis of oocyte defects in control germlines. (A-C)** Graphs showing the total number (n = 290) of germlines analyzed and germlines with specific oocyte defects indicated by graph titles. Percentage of defective germlines is marked at the top of the respective bar. With the exception of oocyte blebbing (5%), no other defects were observed in controls.

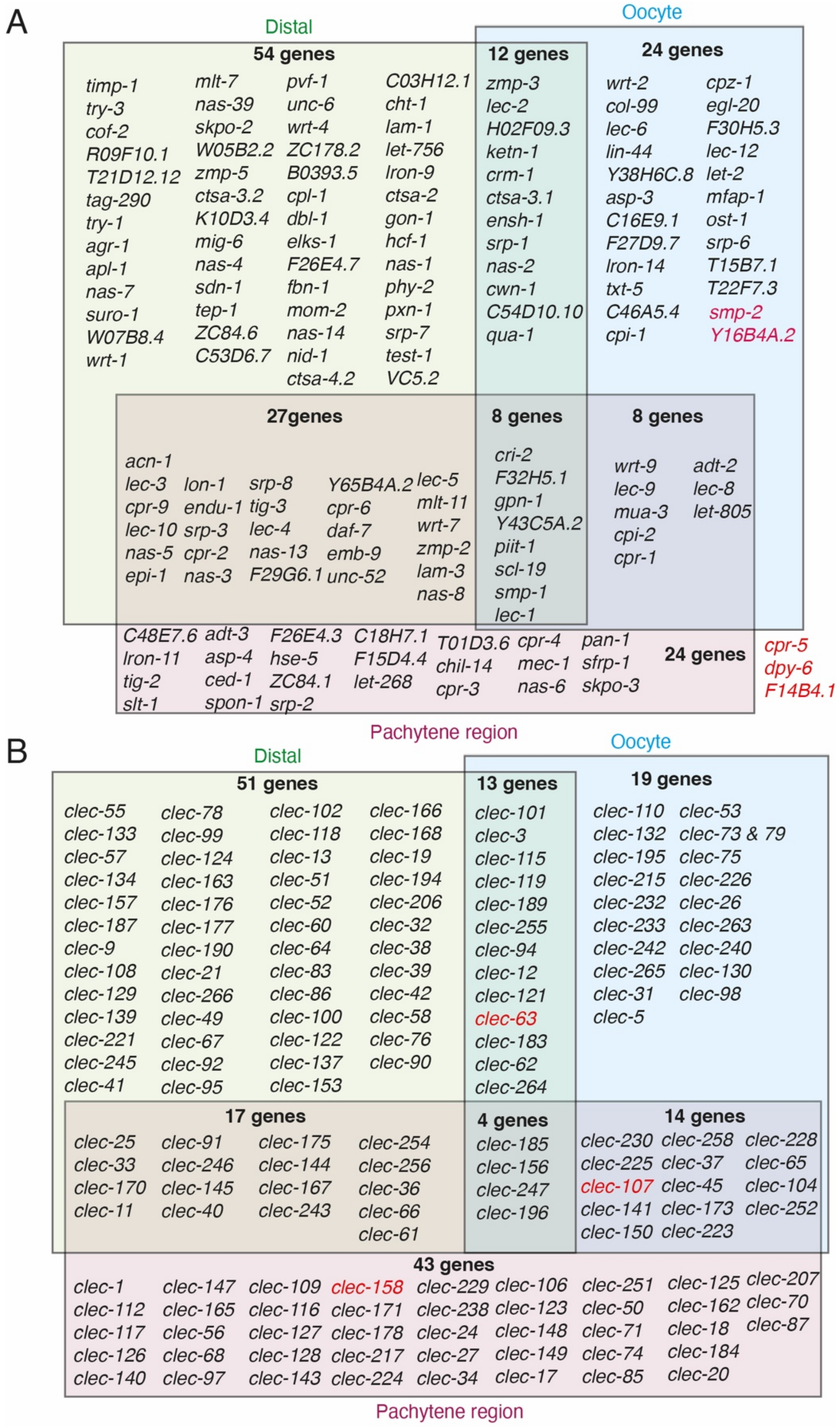


**Fig. S11. Genes showing specific local and global phenotypes. (A)** Non-CLEC genes with phenotypes identified by RNAi. Red letters = germ tube defects **(B)** CLEC genes with phenotypes identified by RNAi. Red letters = germ tube defects

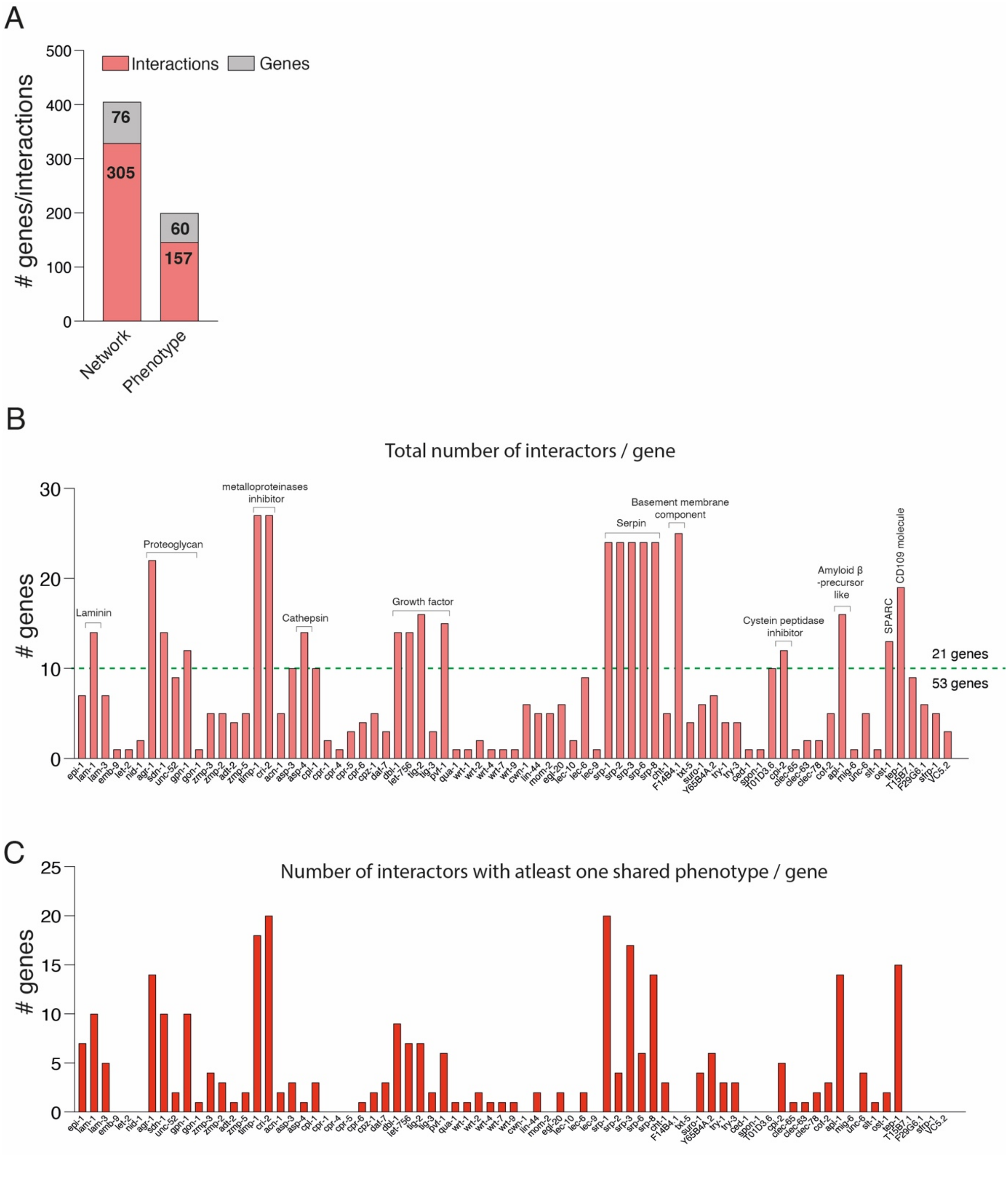


**Fig. S12. Number of gene interactions and shared phenotypes. (A)** 76 genes formed 305 interactions had experimentally verified phenotypes. 60 genes forming 157 interactions had at least one shared phenotype with their interacting partners. **(B)** Total number of interactors per gene. 21 genes showed 10 more interaction partners. **(C)** Number of interactors with at least one shared phenotype with each gene.

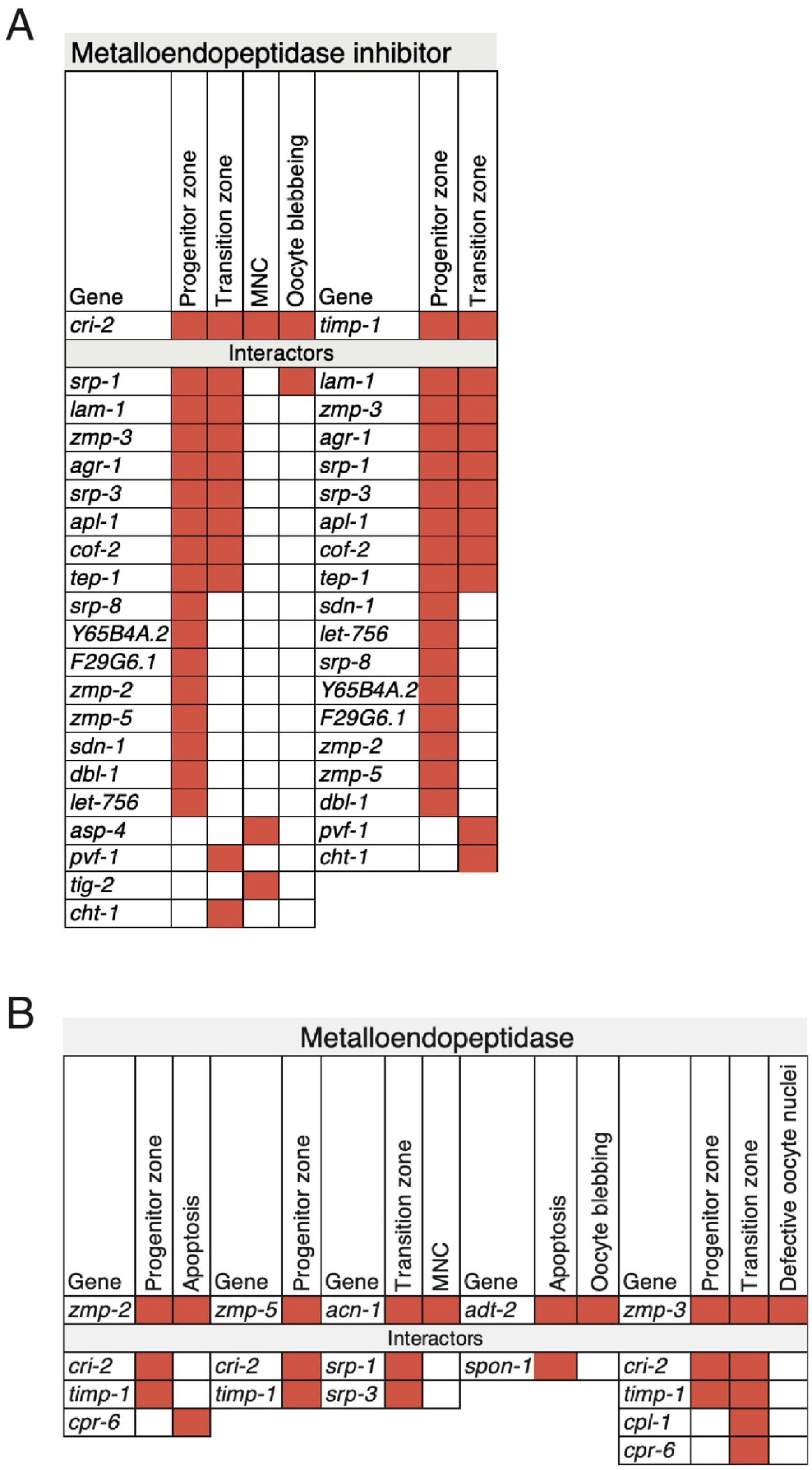


**Fig. S13. Interactions and phenotypes for metalloendopeptidase inhibitors and metalloendopeptidases. (A-B)** Germline phenotypes shared by (A) metalloendopeptidase inhibitors, (B) metalloendopeptidases and their interactors. Red square = identified phenotypes.

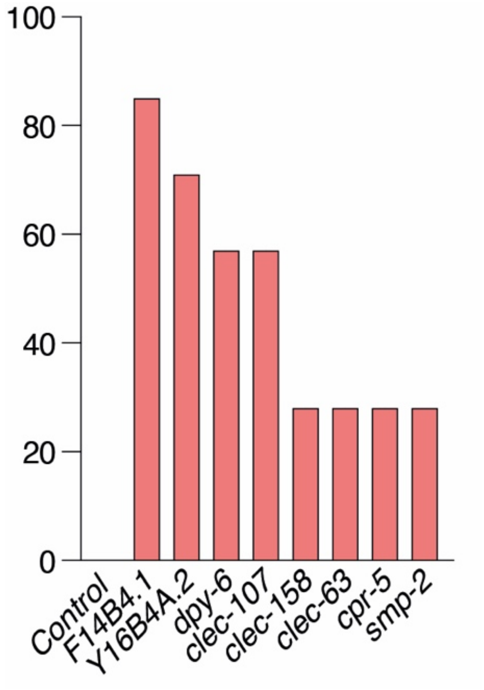


**Fig. S14. Specific matrisome genes showed defects in germ tube morphology**. Graph showing percentage of germlines with germ tube defects following RNAi knockdown.

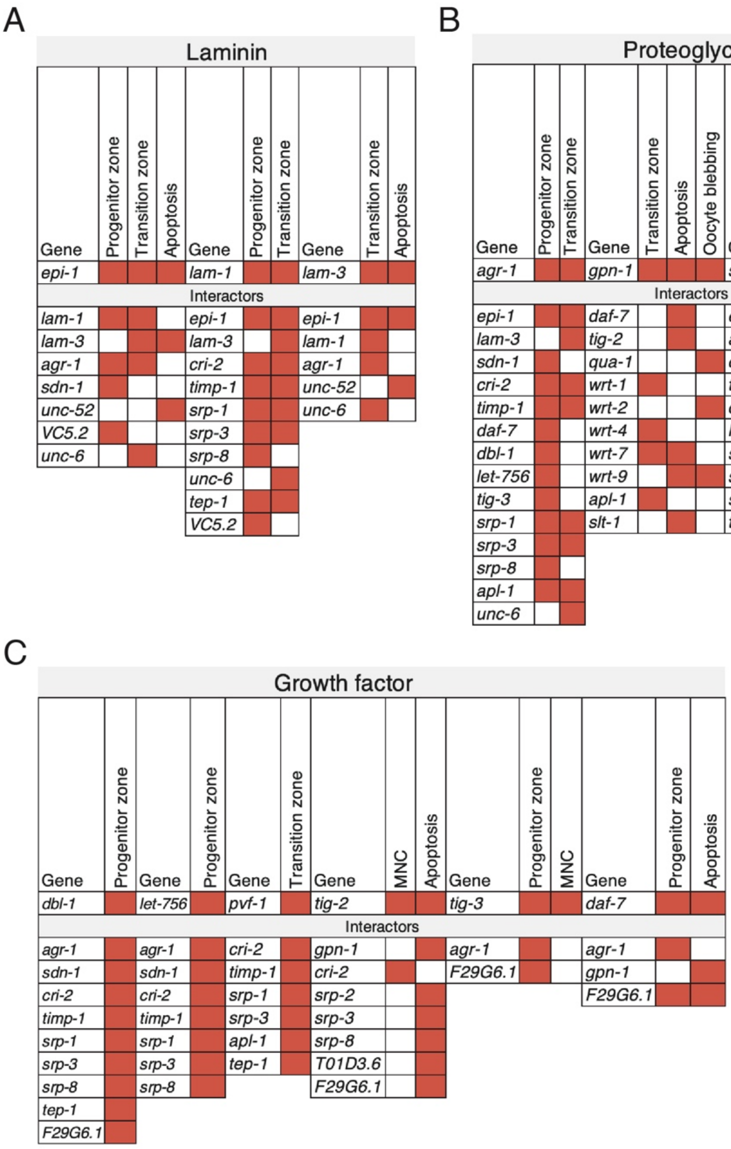


**Fig. S15. Interactions and phenotypes of laminins, proteoglycans, and growth factors.** Interactors of A) laminins, B) proteoglycans, and C) growth factors with shared phenotypes. Red square = identified phenotypes.

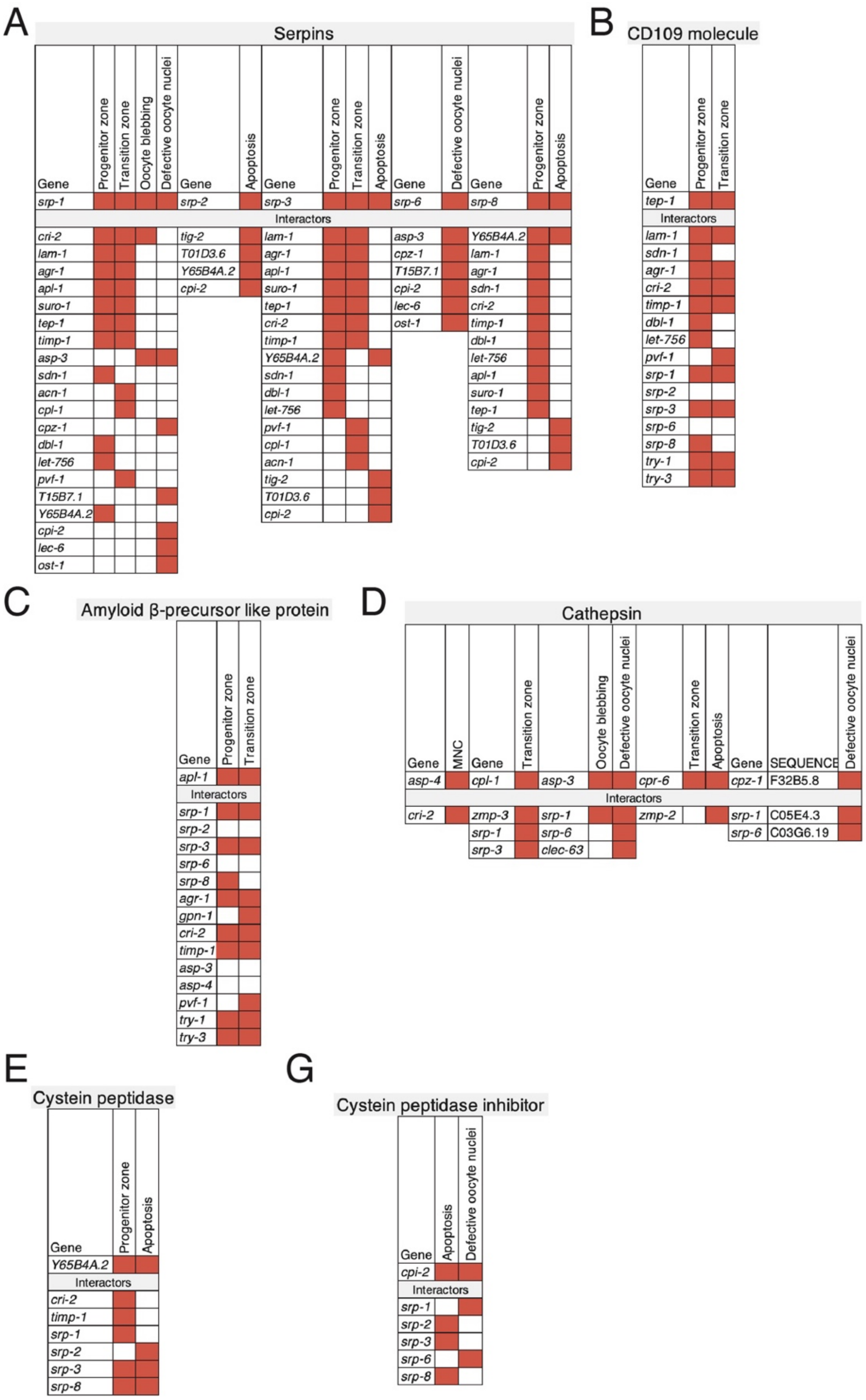


**Fig. S16. Matrisome genes and their interaction partners with shared germline phenotype.** Interactors of **(A)** serpins, **(B)** CD109 molecules, **(C)** amyloid β-precursor like protein, **(D)** cathepsins, **(E)** cysteine peptidases and **(F)** cysteine peptidase inhibitors with shared phenotypes. Red square = identified phenotypes.

A

B

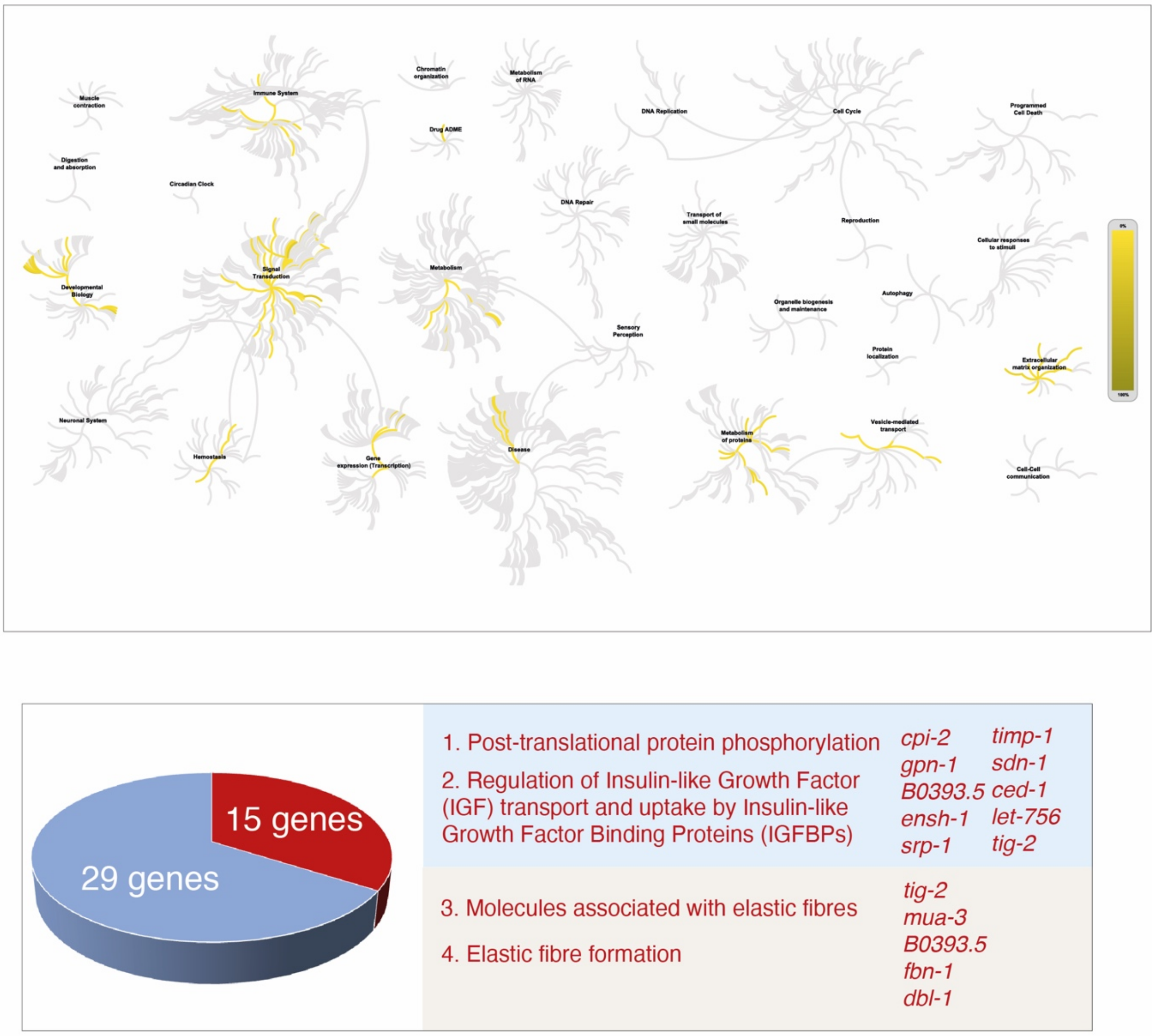


**Fig. S17. Functional pathways of *C. elegans* matrisome. (A)** Overrepresentation analysis of human orthologs of conserved *C. elegans* matrisome genes using the Reactome database showing their roles in 170 signaling pathways. **(B)** 15 genes present in 4 pathways of high confidence (P-value< 0.05).

**Table S1.** Gene knockdowns resulted in altered progenitor zone cell number.

| Gene | Significance |
|---|---|
| agr-1 | **** |
| apl-1 | **** |
| C03H12.1 | **** |
| clec-108 | **** |
| clec-139 | **** |
| clec-157 | **** |
| clec-187 | **** |
| clec-189 | **** |
| clec-245 | **** |
| clec-25 | **** |
| clec-3 | **** |
| clec-33 | **** |
| clec-41 | **** |
| clec-55 | **** |
| clec-57 | **** |
| clec-9 | **** |
| clec-91 | **** |
| clec-95 | **** |
| clec-99 | **** |
| cof-2 | **** |
| cri-2 | **** |
| ctsa-4.2 | **** |
| endu-1 | **** |
| F32H5.1 | **** |
| ketn-1 | **** |
| lec-3 | **** |
| let-756 | **** |
| nas-7 | **** |
| R09F10.1 | **** |
| skpo-2 | **** |
| tag-290 | **** |
| timp-1 | **** |
| try-1 | **** |
| try-3 | **** |
| W05B2.2 | **** |
| W07B8.4 | **** |
| wrt-1 | **** |
| zmp-3 | **** |
| zmp-5 | **** |
| clec-100 | *** |
| clec-119 | *** |
| clec-122 | *** |
| clec-145 | *** |
| clec-170 | *** |
| clec-177 | *** |
| clec-21 | *** |
| clec-246 | *** |
| clec-40 | *** |
| clec-92 | *** |
| clec-94 | *** |
| F29G6.1 | *** |
| H02F09.3 | *** |
| lon-1 | *** |
| lron-9 | *** |
| nas-4 | *** |
| suro-1 | *** |
| T21D12.12 | *** |
| ZC84.6 | *** |
| C53D6.7 | ** |
| clec-101 | ** |
| clec-102 | ** |
| clec-156 | ** |
| clec-167 | ** |
| clec-176 | ** |
| clec-243 | ** |
| clec-247 | ** |
| clec-51 | ** |
| clec-52 | ** |
| clec-63 | ** |

| Gene | Significance |
|---|---|
| clec-78 | ** |
| clec-83 | ** |
| clec-86 | ** |
| crm-1 | ** |
| ctsa-2 | ** |
| daf-7 | ** |
| ensh-1 | ** |
| gon-1 | ** |
| hcf-1 | ** |
| lam-1 | ** |
| lec-1 | ** |
| lec-2 | ** |
| lec-5 | ** |
| mlt-11 | ** |
| qua-1 | ** |
| srp-7 | ** |
| srp-8 | ** |
| tig-3 | ** |
| ZC178.2 | ** |
| zmp-2 | ** |
| clec-11 | * |
| clec-115 | * |
| clec-121 | * |
| clec-12 | * |
| clec-134 | * |
| clec-144 | * |
| clec-168 | * |
| clec-194 | * |
| clec-190 | * |
| clec-221 | * |
| clec-254 | * |
| clec-255 | * |
| clec-264 | * |
| clec-36 | * |
| clec-38 | * |
| clec-39 | * |
| clec-42 | * |
| clec-66 | * |
| clec-76 | * |
| clec-90 | * |
| sdn-1 | * |
| cpr-2 | * |
| cwn-1 | * |
| dbl-1 | * |
| elks-1 | * |
| epi-1 | * |
| lec-10 | * |
| lec-4 | * |
| mig-6 | * |
| nas-2 | * |
| nas-8 | * |
| pxn-1 | * |
| smp-1 | * |
| srp-1 | * |
| srp-3 | * |
| tep-1 | * |
| VC5.2 | * |
| wrt-4 | * |
| wrt-7 | * |
| Y65B4A.2 | * |
| B0393.5 | * |

| | |
|---|---|
| * | p<0.05 |
| ** | p<0.01 |
| *** | p<0.001 |
| **** | p<0.0001 |

**Table S2.** Gene knockdowns resulted in altered transition zone cell number.

| Gene | Significance |
|---|---|
| *nas-39* | **** |
| *cht-1* | **** |
| *mlt-7* | **** |
| *cri-2* | **** |
| *epi-1* | **** |
| *F32H5.1* | **** |
| *T21D12.12* | **** |
| *zmp-3* | **** |
| *ctsa-4.2* | **** |
| *timp-1* | **** |
| *clec-134* | **** |
| *clec-33* | **** |
| *gpn-1* | **** |
| *clec-49* | **** |
| *clec-156* | **** |
| *clec-163* | **** |
| *clec-25* | **** |
| *ctsa-3.2* | *** |
| *piit-1* | *** |
| *K10D3.4* | *** |
| *nas-5* | *** |
| *ctsa-3.1* | *** |
| *try-3* | *** |
| *phy-2* | *** |
| *clec-118* | *** |
| *clec-133* | *** |
| *clec-124* | *** |
| *clec-196* | *** |
| *clec-221* | *** |
| *clec-266* | *** |
| *clec-64* | *** |
| *lec-2* | *** |
| *lec-3* | *** |
| *pvf-1* | *** |
| *clec-67* | *** |
| *clec-170* | *** |
| *clec-61* | *** |
| *lam-3* | *** |
| *clec-11* | *** |
| *clec-32* | *** |
| *acn-1* | ** |
| *mig-6* | ** |
| *tag-290* | ** |
| *tep-1* | ** |
| *suro-1* | ** |
| *clec-115* | ** |
| *clec-129* | ** |
| *clec-13* | ** |
| *clec-190* | ** |
| *R09F10.1* | ** |
| *clec-255* | ** |
| *clec-55* | ** |
| *clec-57* | ** |
| *lec-10* | ** |
| *lec-4* | ** |
| *cof-2* | ** |
| *mom-2* | ** |
| *clec-247* | ** |
| *clec-3* | ** |
| *lam-1* | ** |

| Gene | Significance |
|---|---|
| *nas-1* | ** |
| *nas-3* | ** |
| *unc-6* | ** |
| *clec-60* | ** |
| *unc-52* | * |
| *nid-1* | * |
| *scl-19* | * |
| *C54D10.10* | * |
| *nas-14* | * |
| *cpr-9* | * |
| *nas-13* | * |
| *cpl-1* | * |
| *F26E4.7* | * |
| *cpr-6* | * |
| *try-1* | * |
| *nas-7* | * |
| *clec-102* | * |
| *clec-121* | * |
| *clec-12* | * |
| *clec-177* | * |
| *clec-176* | * |
| *clec-157* | * |
| *clec-19* | * |
| *clec-183* | * |
| *ketn-1* | * |
| *clec-254* | * |
| *agr-1* | * |
| *W07B8.4* | * |
| *clec-9* | * |
| *H02F09.3* | * |
| *fbn-1* | * |
| *wrt-1* | * |
| *wrt-4* | * |
| *wrt-7* | * |
| *Y43C5A.2* | * |
| *clec-144* | * |
| *clec-206* | * |
| *clec-58* | * |
| *clec-62* | * |
| *clec-175* | * |
| *srp-3* | * |
| *clec-153* | * |
| *apl-1* | * |
| *cpr-2* | * |
| *clec-185* | * |
| *clec-256* | * |
| *test-1* | * |
| *emb-9* | * |
| *clec-166* | * |
| *clec-187* | * |
| *clec-137* | * |

| | |
|---|---|
| * | p<0.05 |
| ** | p<0.01 |
| *** | p<0.001 |
| **** | p<0.0001 |

**Table S3.** Genes knockdowns resulted in significant changes in apoptosis in the germline.

| Non-clecs | |
|---|---|
| **Gene** | **Significance** |
| *adt-2* | * |
| *lec-4* | * |
| *him-4* | * |
| *cpi-2* | ** |
| *lec-5* | * |
| *cpr-1* | ** |
| *F29G6.1* | * |
| *lec-1* | * |
| *pan-1* | * |
| *srp-2* | * |
| *C48E7.6* | **** |
| *srp-3* | ** |
| *cpr-4* | * |
| *emb-9* | * |
| *endu-1* | * |
| *lec-8* | * |
| *cpr-6* | * |
| *daf-7* | * |
| *let-268* | ** |
| *unc-52* | * |
| *zmp-2* | * |
| *cpr-3* | * |
| *spon-1* | * |
| *lec-3* | * |
| *piit-1* | * |
| *skpo-3* | * |
| *wrt-9* | * |
| *mlt-11* | * |
| *gpn-1* | *** |
| *chil-14* | * |
| *nas-6* | * |
| *wrt-7* | * |
| *tig-2* | * |
| *let-805* | * |
| *mec-1* | * |
| *sfrp-1* | * |
| *slt-1* | * |
| *lam-3* | ** |
| *nas-3* | *** |
| *Y43C5A.2* | **** |
| *C18H7.1* | ** |
| *F15D4.4* | ** |
| *nas-8* | ** |
| *T10D3.6* | ** |
| *Y65B4A.2* | ** |
| *cpr-9* | *** |
| *srp-8* | *** |
| *epi-1* | * |
| *F26E4.3* | *** |
| *nas-5* | **** |
| *lron-11* | **** |

| Clecs | |
|---|---|
| **Gene** | **Significance** |
| *clec-243* | ** |
| *clec-162* | * |
| *clec-65* | ** |
| *clec-125* | * |
| *clec-91* | * |
| *clec-87* | * |
| *clec-233* | * |
| *clec-251* | ** |
| *clec-104* | * |
| *clec-11* | * |
| *clec-167* | * |
| *clec-18* | * |
| *clec-184* | * |
| *clec-20* | * |
| *clec-207* | * |
| *clec-252* | * |
| *clec-33* | * |
| *clec-40* | * |
| *clec-70* | * |
| *clec-123* | ** |
| *clec-106* | ** |
| *clec-256* | ** |
| *clec-143* | *** |
| *clec-148* | ** |
| *clec-149* | ** |
| *clec-17* | ** |
| *clec-173* | ** |
| *clec-223* | ** |
| *clec-228* | ** |
| *clec-230* | ** |
| *clec-36* | ** |
| *clec-50* | ** |
| *clec-71* | ** |
| *clec-74* | ** |
| *clec-85* | ** |
| *clec-254* | *** |
| *clec-117* | **** |
| *clec-112* | **** |
| *clec-126* | **** |
| *clec-130* | **** |
| *clec-140* | **** |
| *clec-147* | **** |
| *clec-165* | **** |
| *clec-175* | **** |
| *clec-225* | **** |
| *clec-56* | **** |
| *clec-66* | **** |
| *clec-97* | **** |

Increase in apoptosis

[illegible]crease in apoptosis

| | |
|---|---|
| * | p<0.05 |
| ** | p<0.01 |
| *** | p<0.001 |
| **** | p<0.0001 |
